# The Wnt segment polarity pathway and TMED2 protein may interact via a lectin- and decoy-type mechanism

**DOI:** 10.1101/056531

**Authors:** Klaus Fiedler

## Abstract

The Secl4-like protein (280-385 GOLD-domain) in this study scores highly with the carbohydrate-binding module (CBM) of the RetS domain from *Pseudomonas aeruginosa.* The thereupon modeled *Cricetulus griseus* p24-GOLD domain of p24, a member of intra-Golgi cargo receptors, is shown to interact with Wnt8 (wingless 8) of *Xenopus laevis* with a ΔG=-18.3 kcal/mol. Lower ranked models listed a smaller ΔG (PDBePISA) and energy of DelPhi interaction. Complex/hybrid N-glycans provide increasing energy of binding up to −7.1 kcal/mol to simulated p24-GOLD-ligand interaction. It is likely, that Wnt proteins and p24 cargo-receptors interact analogously to Wnt-Frizzled and that Wnt transport may involve early lectin binding. The possibly promiscuous interaction of p24-GOLD with ligands, including collagen, may shed light on cargo-receptor mediated traffic.

**Highlights:**

- p24s are usually described as type I transmembrane proteins containing a GOLD domain
- p24-GOLD can be modeled based on the Sec14-like protein with X-ray structure quality
- A concave patch on the p24 surface binds to diverse ligands
- N-glycans may provide increased binding energy in p24-cargo interaction
- Wnt binding to TMED2 may quench signaling similar to a decoy-type mechanism

## Introduction

Galectins and L-type lectins have previously been shown to be involved in apical delivery in MDCK (Madin-Darby Canine Kidney) and gastrointestinal tumor cells (1–3). A clustering of secreted proteins, galectins and Forssman-antigen, a globoside, possibly mediates apical sorting (4) and integral-membrane adaptors, such as VIP36, may trigger coat-mediated delivery (5–11). VIP36 for example binds to high-mannose N-glycans (12). Although the principles of sorting may be established, a difference of species and expression of glycosyltransferases has not led to a consistent description of apical delivery in higher eukaryotes (see *in vivo* expression (13)). An urgent need exists for comparative analysis of glycosyltransferases as well as lectins that may interact with glycans of secreted and Golgi-resident enzymes to reconcile results that have been obtained in MDCK and other cells with human biology.

The TMED (transmembrane emp24 domain containing) family of cargo receptors have recently been found in Golgi-derived vesicles in CHO (chinese hamster ovary) cells and in *S. cerevisiae* and are described as oligomeric, dimeric and monomeric integral membrane proteins (14–20). Emp24-gp25L-p24 domains of p24/TMED proteins and p24-like proteins are found in 285 proteins in Chordata, the function of these proteins is not entirely understood and could involve insulin secretion in pancreatic β-cells (21). Early embryonic lethality had been found for gene-deleted TMED2 (p24ß1) mice which has led to the suggestion, that despite lack of evidence for *gross* growth defects in *S. cerevisiae,* early development in mammals, and possibly unique growth factors, are activated by p24 function (22, 23). These analyses on the *S. cerevisiae* protein showed that the enzymatic chain of N-glycosylation as such was not affected in gene-deleted yeast strains (19).

The question has been raised whether in eukaryotes early transport steps in the secretory pathway would involve sorting of GPI-anchored proteins recognized by TMED cargo receptors (19, 24). GPI-anchored proteins are attached to the cell surface and are sorted to the apical plasma membrane in polarized cells as shown for example in MDCK and other cells (10, 11). In MDCK cells it has been elucidated that the apical transport involves both, cholesterol-mediated rafts and N-glycosylation for GPI-anchored proteins. In FRT (Fischer rat thyroid) cells both cell surfaces, the basal and apical side receive outgoing Golgi traffic and some sorted GPI-anchored proteins and these routes were shown to be independent of cholesterol (25, 26).

A similarity of Sec14-like proteins to p24 proteins had previously been suggested based on primary sequence comparison (27). In the present analysis I have compared the p24 protein TMED2 with PDB structure data and find a similarity of p24 to CBMs (28, 29) and to Sec14-like protein (30, 31) that both carry modules structurally homologous to the p24-GOLD (Golgi-dynamics) domain (27). Surprisingly, p24-GOLD interacts with glycans with intermediate affinity as shown in chemical docking. Also Wnt8 (32, 33), Wnt8(C) being involved in epithelial mesenchymal transition (EMT)(34, 35), interacts in a high affinity configuration with p24-GOLD, suggesting that the type of lectins involved in apical sorting in epithelial cells may be more diverse than previously assumed.

For a full account of the complexity of protein sorting and of cell polarity and cancer the reader is referred to (35–37). Links of protein sorting and EMT are listed in these reviews.

## Materials and Methods

Remote homology tracking and structure prediction was achieved with HHM (Hidden Markov Model) comparison that incorporates the PSIPRED predicted secondary structure score and relates these to actual or predicted DSSP values for structures of database entries (38) (http://toolkit.tuebingen.mpg.de/hhpred). The search was carried through on February 19, 2015. The probability of a template to be a true positive is calculated from the distribution of amino acid residues in the alignments using the secondary structure score. The secondary structure predicted search with local alignment was used to list all output. The alignment of p24 included residues 17-111 of the full length protein from *Cricetulus griseus.* MODELLER was used to generate a first GOLD-domain model of p24 (http://salilab.org/modeller)(39). Residues 16-109 corresponded to a compact module including the mature N-terminus of the protein up to the C-terminal coiled-coil region and was modeled using a template from 4UYB (Sec14-like protein 3)(31). The validity of the structure was confirmed by comparison to PDB entries scoring absolute values of quality QMean (40). These included Cß interaction, all-atom interaction, solvation, torsion, predicted and calculated secondary structure- and predicted-relative and relative solvent accessibility-agreement resulting in Z and Qmean values.

The ClusPro modeling queue was used for structural modeling of protein-protein interactions (41) with default settings (without structure modification or PDB masking) of the chop24a GOLD-domain 16-109 and residues 32-338 of XWnt8 from *Xenopus laevis* (32). Further family members of each protein were searched and visualized with ENSEMBL (42). MUSCLE 3.7 was from R. Edgar (43). The sequence logo is from Prosite (http://prosite.expasy.org). SeaView 4.4.2 was from M.G. Gouy (44). Clustal Omega was used for further multiple sequence alignments with default values except iterations that were set to 5 (45). In phylogenetic tree generation the neighbor-joining algorithm was used with default values (no multiple substitution correction; include gaps) (46–48). PatchDock was used for hard sphere docking of collagen and other proteins that were directly compared, results were usually refined by FireDock (49) unless otherwise indicated. To validate different docking procedures, PDBePISA v1.5.1 (50, 51) interface determination was used. Proteins were selected from PDB and UniProt by logical searches. ENSEMBL was used for initial selection of transmembrane p24/TMED proteins, sequences that did not contain a predicted transmembrane domain at the C-terminus with ~ < 51 residue cytoplasmic domain were not listed as TMED members. SignalP-4.1 was from http://www.cbs.dtu.dk/services/SignalP-4.1 (52).

Docking of glycans was carried through with the conformer of p24a (ClusPro modified) of residues 16-109 (GOLD-domain) and carbohydrates obtained from Dr. Woods’ website (http://www.ccrc.uga.edu) (with multiple conformers) or the Glycosciences website (http://www.glycosciences.de) and the PyRx modeling queue Version 0.8. The carbohydrate collection was selected and filtered for uniqueness against a laboratory collection of glycosphingolipid (GSL)-glycan headgroups that was built using the chemical descriptor-library of LIPIDMAPS (http://www.lipidmaps.org). Pdb files obtained were tested for atomic bonds and partial atomic models were completed using structure viewer programs. Prehydrogenized models were energy-minimized using MarvinView 6.1.3 (Dreiding force field) (ChemAxon Ltd.) and files were processed by Chimera 1.8.1 (http://www.cgl.ucsf.edu/chimera) for import to PyRx docking. The AutoDock VINA (53) implementation of PyRx from S. Dallakyan (http://pyrx.scripps.edu) was utilized with the grid size as indicated in single experiments. The algorithm installs OpenBabel (54) and a uff (united force field) for energy minimization, conjugate gradients with 200 steps and a cut-off for energy minimization of 0.1. Partial charges were added to receptors using PyBabel (MGL Tools; http://mgltools.scripps.edu). No limits to torsions were allowed. Single CPU time was up to 250 hours for longest/branched ligands in exhaustiveness 11. Sqlite data were analyzed using SQLite (Hipp, D. R.).

Structural comparisons were carried through with PDBeFold v.2.59 (searched 26. Feb. 2015). DelPhi was used to evaluate electrostatic interactions of protein complexes (http://compbio.clemson.edu/delphi)(55) with submission settings: Interior dielectric constant 2, exterior dielectric constant 80, %fill 60, grid scale 1.2, salt concentration 0 or 140 mM, probe radius 1.4, non-linear and with calculations in Excel spreadsheets (Microsoft Excel 2007). Molecular optimization was carried through with Ascalaph Version 1.8.88 Molecular Modeling. The OPLS and implicit water (Sheffield) was chosen for hybrid Lui Storey and conjugate descent optimization at step 0.00001, gradient 0.000001, function 0.000001 and default control with 27 iterations with an RMSD of 0.123 (protein). The energy of binding increased by 6.3%.

H-bonding was determined with ViewDock and with tolerances 0.4 Å, 20° (56) or 0.8 Å, 30° similar to calculations previously applied (51), or as secondarily found in molecular mechanics with the former determination (56). Annotation of carbohydrates was from Dr. Woods’ website and from glycosciences.de. Glycosylation of proteins was carried through with pdb modifications *in silico* by GlyProt (57). Glycan compound drawing was done with GlycoWorkbench (58). Local computing was carried through with i5 processor, Windows and Ubuntu with Linux 3.13.0-46 system. Chimera 1.8.1 was used for further calculations and hydrophobic/electrostatic surface presentations using default values.

## Results

### Structural Searches with p24-GOLD

The Sec14-like protein supernatant protein factor (30) (SPF) has previously been shown to be similar to Sec14 and to include a jelly-roll domain at its C-terminus that seemed unrelated to its lipid binding activities. It has been proposed that it may bind to further Golgi proteins but the similarity to viral proteins or receptor binding domains had not provided a clear indication of its function. I have tested the similarity of the C-terminal GOLD-domain (which is present in 3’312 proteins (Uniprot_2014_6) and 2 structures (SCOP 1.75)) to structure entries of the current PDB database. Not only the CRAL/TRIO-domain (30) but also the Golgi localization three-helix coil (Fig. 1) were not analyzed in this analysis by PDBeFold. The GOLD-domain is suggested to fold into an independent structural domain and simple structural modelling seemed thus feasible (27). The structure alignment showed a high Q-score for the collagenase collagen-binding domain 0.42 of *Clostridium histolyticum* which had previously been shown to increase processivity of the enzyme carrying two collagen-binding domains (59) (Suppl. Table S2). In addition to endo-1,4-ß-galactosidase further carbohydrate-binding modules (60), named CBMs, were showing high Q-scores of 0.3 and above. Galectins (61) are also listed in the structural similarity search but may not be equated with a jelly-roll fold of the typical 4+4 antiparallel ß sheets of p24 and other GOLD-domains (30). The whole protein and concave face of the jelly-roll fold of CHOp24 was analyzed further for N-glycan binding (see Suppl. Fig. S1, S2).

**Figure 1.**
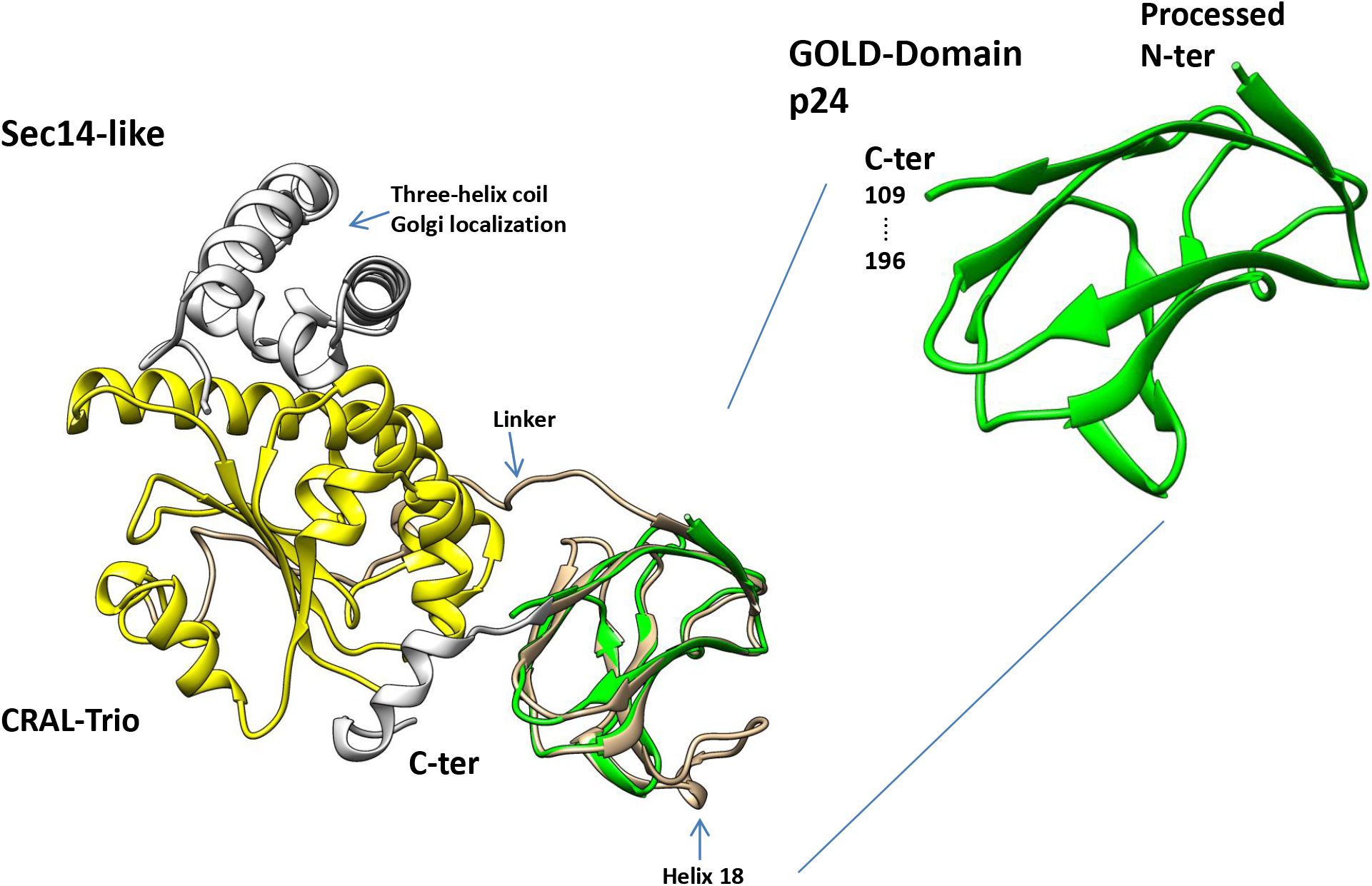
The Sec14-like protein and similarity to the p24-GOLD-domain. Sec14-like protein 3 includes two distinct domains, a CRAL-TRIO and GOLD domain. CRAL-TRIO (yellow) binds to lipids (yeast Sec14) and tocopherol (human tocopherol transfer-protein); the orthologue Sec14-like protein 2 was crystallized and found to display a closed lipid-exchange loop (residue 196-215) which shields the hydrophobic binding cavity. A C-terminal helix (Helix 20 in the Sec14-like protein 3 / 4UYB) (silver) might stabilize the lipid-exchange loop but is otherwise unrelated to Sec14-like protein GOLD-domain (gold) cargo-receptor activity. The p24 GOLD-domain (green) models with crystallographic-like quality from residue 16 to 109. This includes the mature, processed N-terminus of the protein following signal cleavage after translocation across the endoplasmic reticulum membrane. The eight-stranded fold of the GOLD domains is similar to unique bacterial collagen- and frequent carbohydrate-binding modules. A linker attaches the Sec14-like GOLD domains to the CRAL-TRIO. Helix 18 of the Sec14-like protein 3 is indicated and shields one side of a concave face oriented opposite to the Golgi-localization structurally described three-helix coil.

### Structural Model of p24-GOLD

In Figure 2 I show the details of the p24 *(Cricetulus griseus)* model with the surface charge distribution of the protein. It was modeled with a Sec14-like protein template (Suppl. Fig. S1). The N-terminus of the protein can be found at 12 o’clock and is followed by 8 ß-strands colored from dark blue, light blue, green and yellow to red in the backbone presentation. A belt of positive charges runs from left to right in the on face view (middle panel) and from top to the bottom on the backside. The N-terminus corresponds to the processed, mature sequence following signal-peptide cleavage after ER (endoplasmic reticulum)-translocation. It is followed by a sequence with coiled-coil structural propensity (14) that had been shown to mediate oligomerization in different p24 family members (18, 62). The membrane span shows an activity in sphingomyelin binding (63) and is attached to a cytoplasmic tail sequence that distributes the protein in the early secretory pathway and presumably Golgi-apparatus by COPI-and COPII-interactions (17,64,65)(conserved residue in Fig. 4B). The model was then applied to PyRx docking with a library of 56 carbohydrates and showed a presumed cut-off value of −6.5 kcal/mol in unrelated docking procedures with a homologous protein (see Fig. 4; no particular similarity of TMED2 and 10). To this end, a p23 model was created with *Mus musculus*

**Figure 2.**
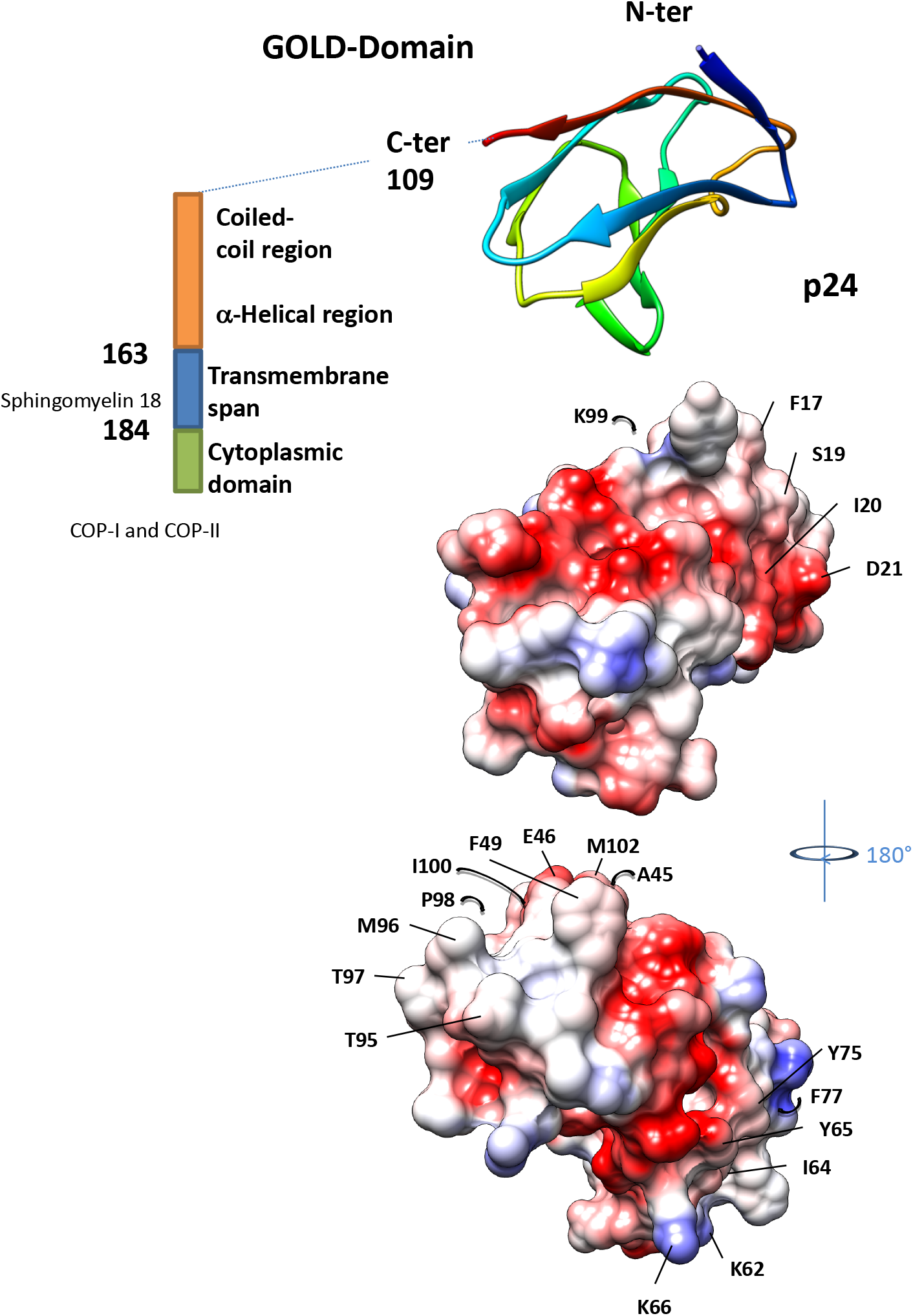
Modeled p24 GOLD-domain. A coiled-coil stretch was found distal to the identified GOLD-domain in sequence analyses (14) which is known to mediate dimerization and oligomerization with other p24 family members. The membrane span includes targeting signals that mediate transit in the secretory pathway and binds to sphingomyelin. The cytoplasmic domain mediates exit via COPII-interaction and Golgi traffic via COPI-binding. Typical and untypical KKXX-signals for “ER retention” are found in the C-terminal stretch of p24 members. The electrostatic surface presentation is shown in Coulombic coloring (settings: 298 K, red, white, blue refer to: dark read −41.9 kJ/(mol x e), dark blue 41.9 kJ/(mol x e); dielectric constant 4, distance from surface 1.4 Å and distance dependence of the dielectric) with identical (top panel) and rotated orientation. The residues identified interacting with Wnt8 are labeled.

TMED10 which due to present lack of modeling quality showed a presumed deep random binding site with an affinity of −6.4 kcal/mol with the top scoring carbohydrate. In stark contrast, p24 was modeled with the carbohydrate attached to a shallow binding groove on the surface facing the viewer in Figure 3A. The ranking of docked carbohydrates shows top-scores with hybrid and complex glycans. One glycan contains a core fucose added, another shows a GlcNAcT-I (N-acetylglucosamine-transferase I) extension with further processing by galactosyl- and sialyltransferase, typical trans-Golgi enzymes. The highly scoring hybrid N-glycan of −7.1 kcal/mol affinity is a product of Golgi-Man II (Golgi-mannosidase II) and is generally found processed to this N-glycan in the medial-Golgi apparatus (66) (see Fig. 5A and Table S3A).

**Figure 3.**
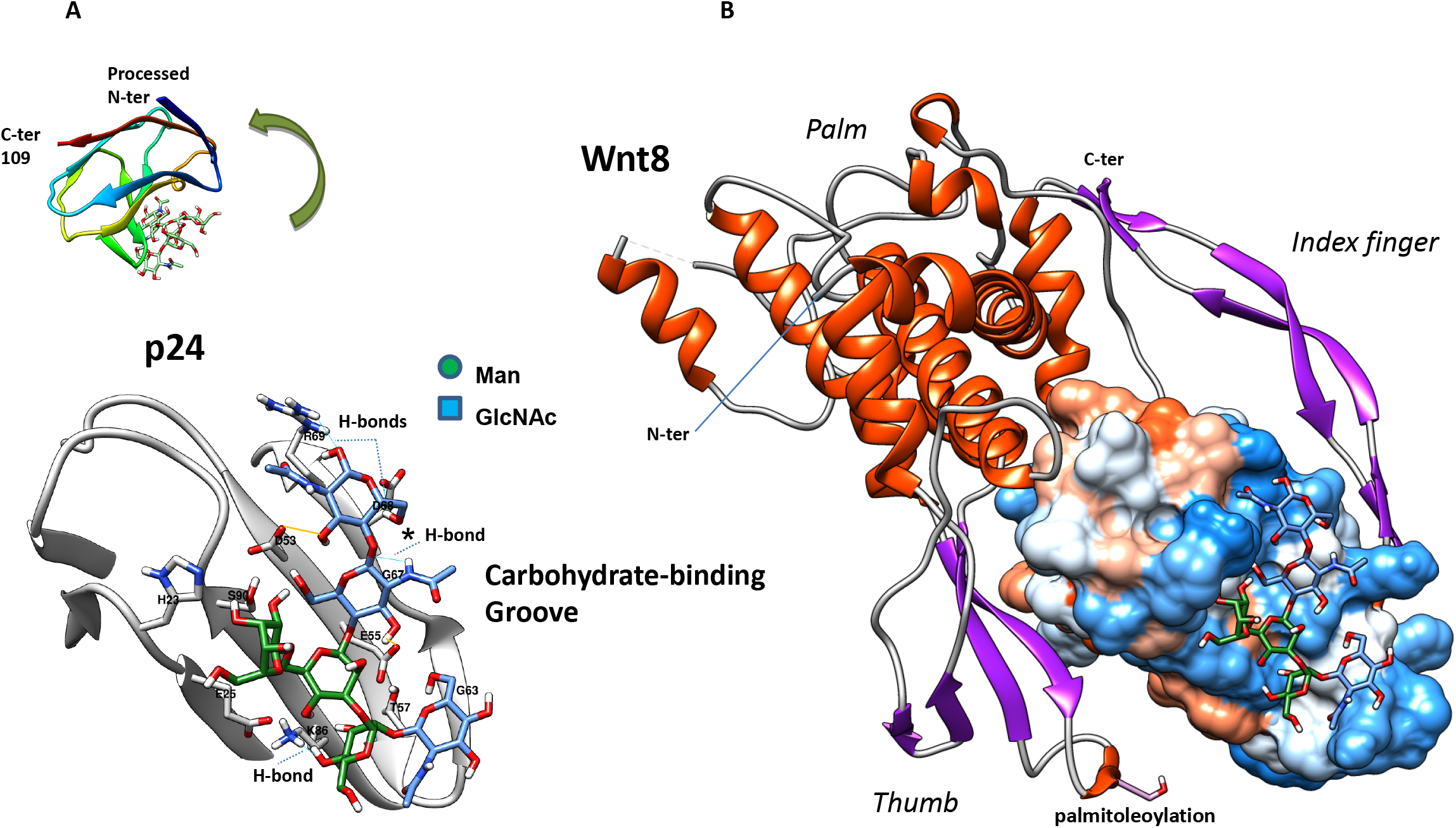
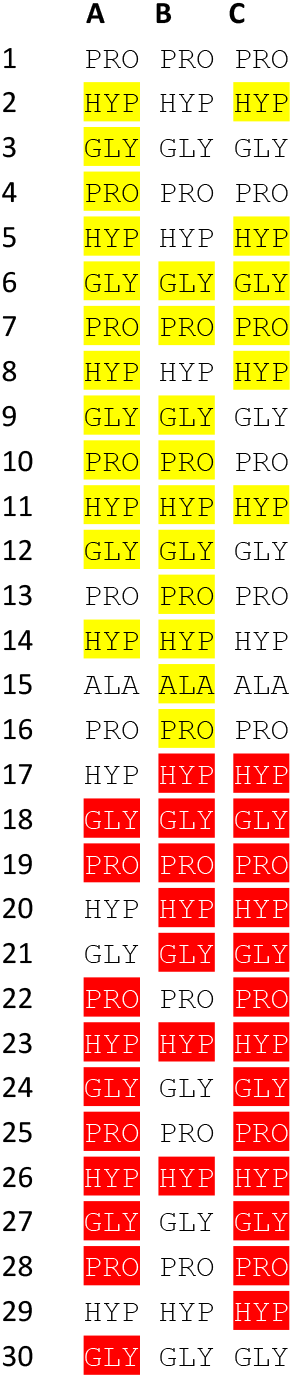
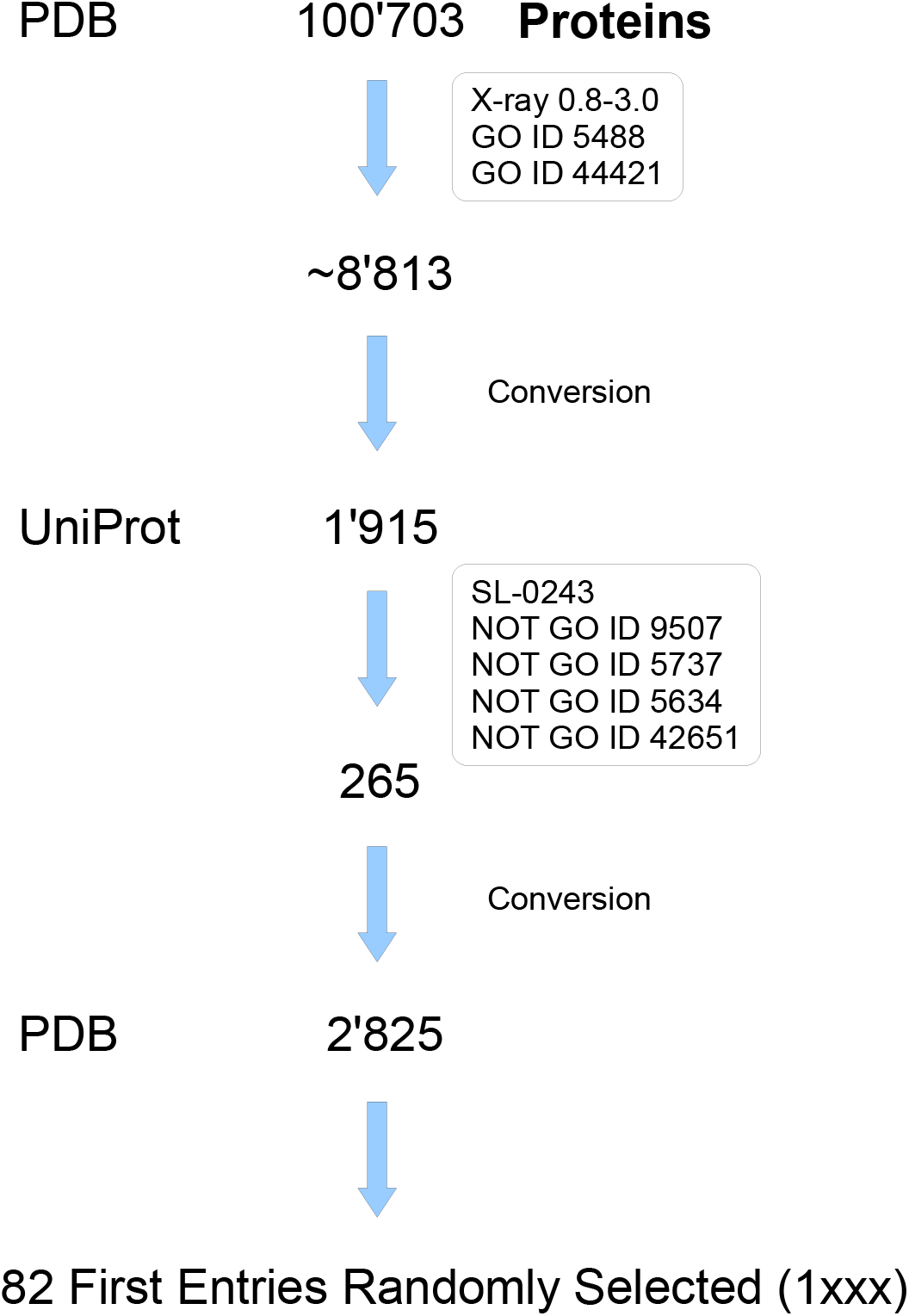
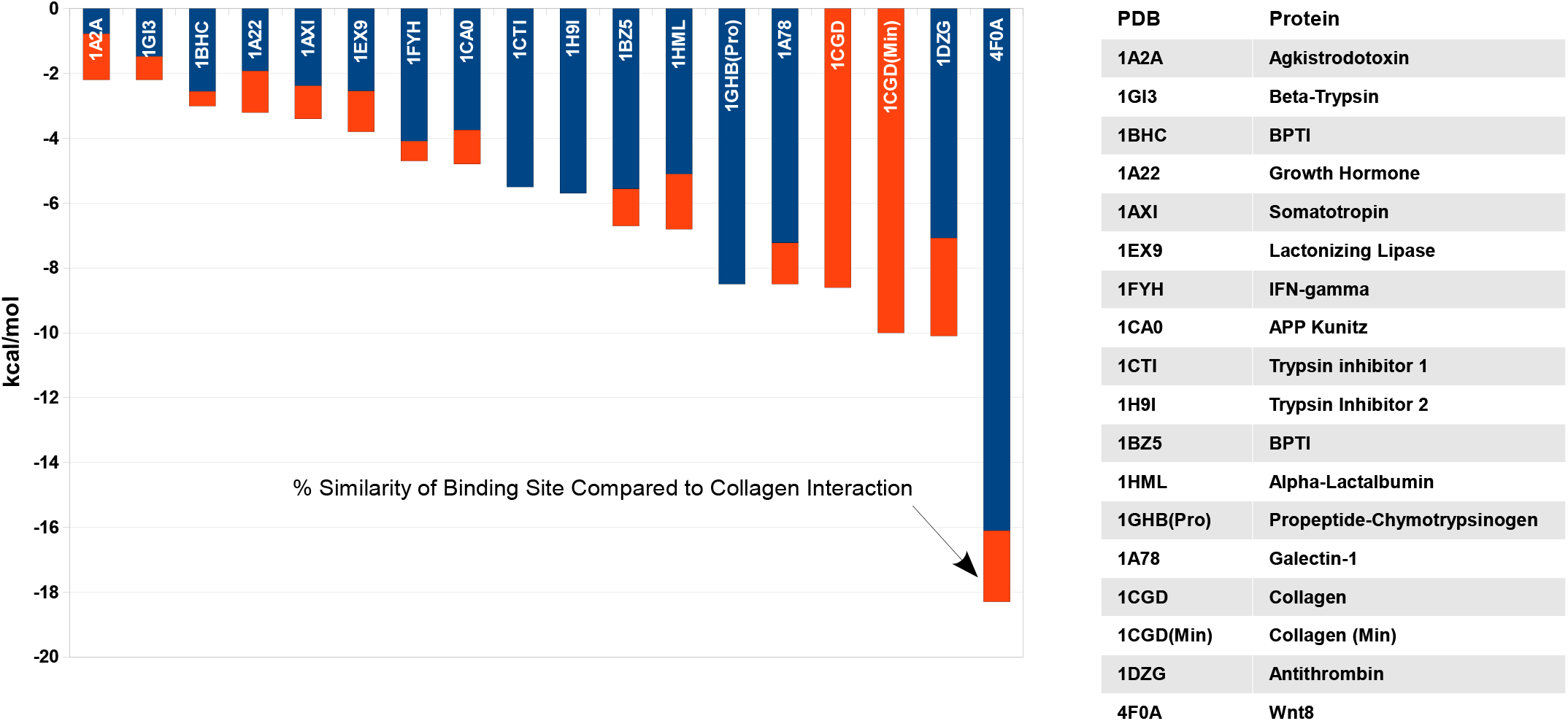
Glycan docking and protein interaction. **A** A carbohydrate binding groove is identified in docked TMED2 from CHO with intermediate affinity (−7.1 kcal/mol) which is similar to other lectins with affinity of approx. 10^4^ M^−1^ to N-glycans (not shown). The hybrid glycan (GlcNAc-core-Man3-GlcNAc) is colored in IUPAC code. Residues on the concave face of docked p24 are labeled within the distance of relaxed hydrogen-bonding. The 4 H-bonds are indicated in blue. Residue Gly67 lost an H-bond (labeled with a *) in molecular mechanics simulation, additional H-bonds were gained in this hybrid simulation with contacts to Asp53 and one H-bond to Glu55. **B** Interaction of p24 cargo receptors with Wnt8 were studied in ClusPro-docking and identified a binding site of the GOLD-domain overlapping with the Fz8 cysteine-rich domain interface at the index finger and a thumb-binding site that was shifted towards the Wnt8 palm (RMSD 0.406 of the docked p24 protein). This is distant from the Wnt8 palmitoleoylation at Ser187. The palmitoleoyl group attaches to Fz8, stabilizes interaction and mediates Fz8 growth factor activity. Although thumb binding is not identical the p24 family proteins may act as a decoy in Wnt interaction. Wnt8 is shown in secondary structure coloring (α-helix, red; sheet, magenta), the p24 surface is colored by hydrophobicity (red, hydrophobic; blue, hydrophilic). **C** Hard-sphere docking was used to analyze the hypothetical interaction of collagens with different Cys-rotamers of TMED2 since TMP21 (TMED10) was found as a chondrocyte-associated antigen. 1CGD was previously used in docking and showed preferred interaction of the N-terminus with TMED2 if in its disulfide (colored yellow) and interaction of the C-terminus with TMED2 if in its free Cys-rotamerization (colored red). **D** PDB was searched for structures according to the scheme presented including terms for intra- and extracellular localization (see UniProt). 2’825 remaining entries were randomized and 82 entries from the 1XXX series were alphanumerically selected. **E** Docking of 82 proteins from the PDB archive randomly selected (FireDock). Collagen is included for comparison in non-optimized (PatchDock) and optimized TMED2 interaction (1CGD(Min)). All structural solutions with terminal collisions were deselected from the ranking procedures. Next to 1CGD antithrombin is highly scoring (1DZG), other 14 are listed. Similarity of the binding site is indicated in orange (% 4 Å distance) by determining the fraction of residues of each interface present in the collagen-docked complex. Wnt8 interaction could likely not successfully be analyzed for hard-sphere interaction since the interface is energetically determined (ClusPro result is indicated).

**Figure 4.**
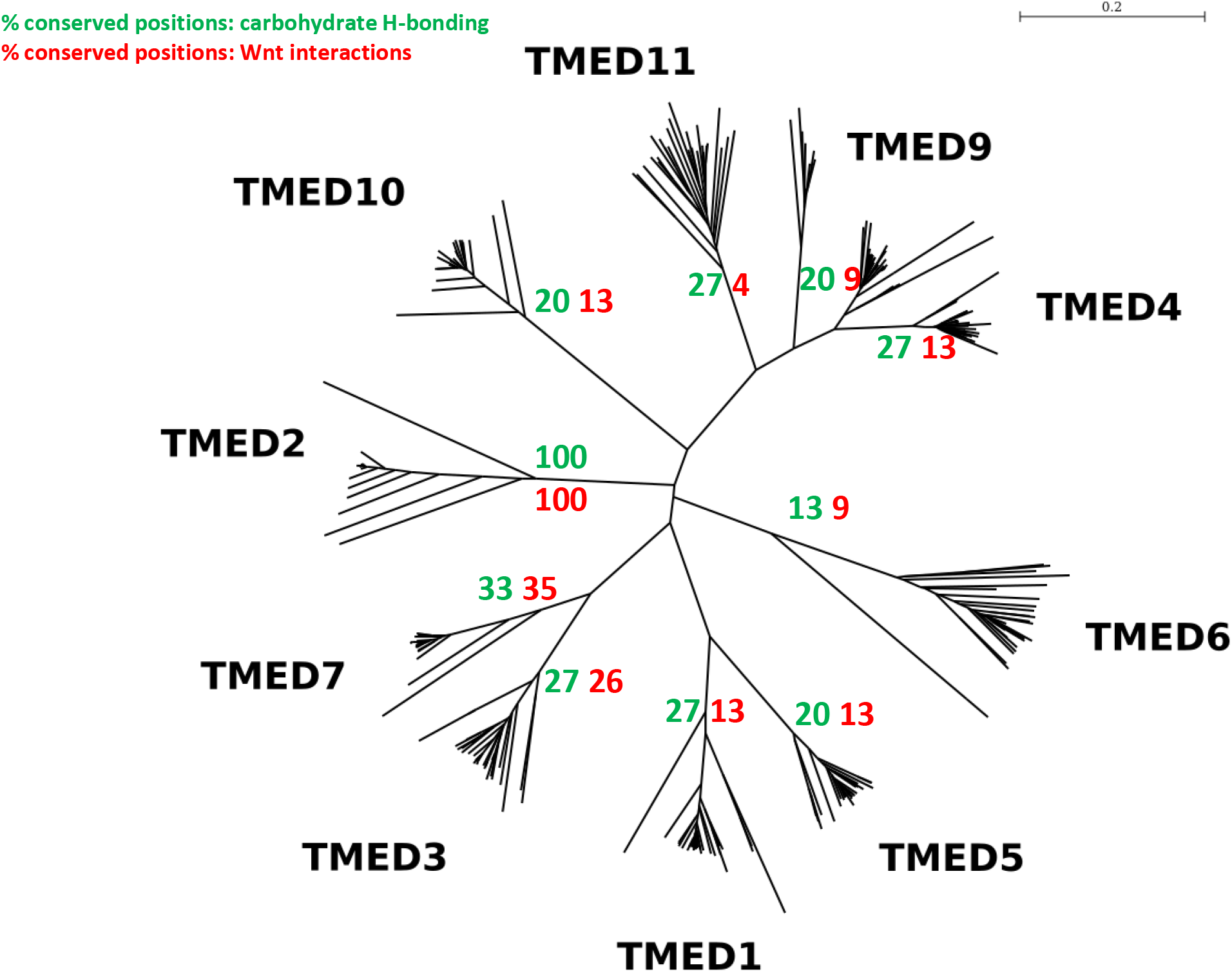
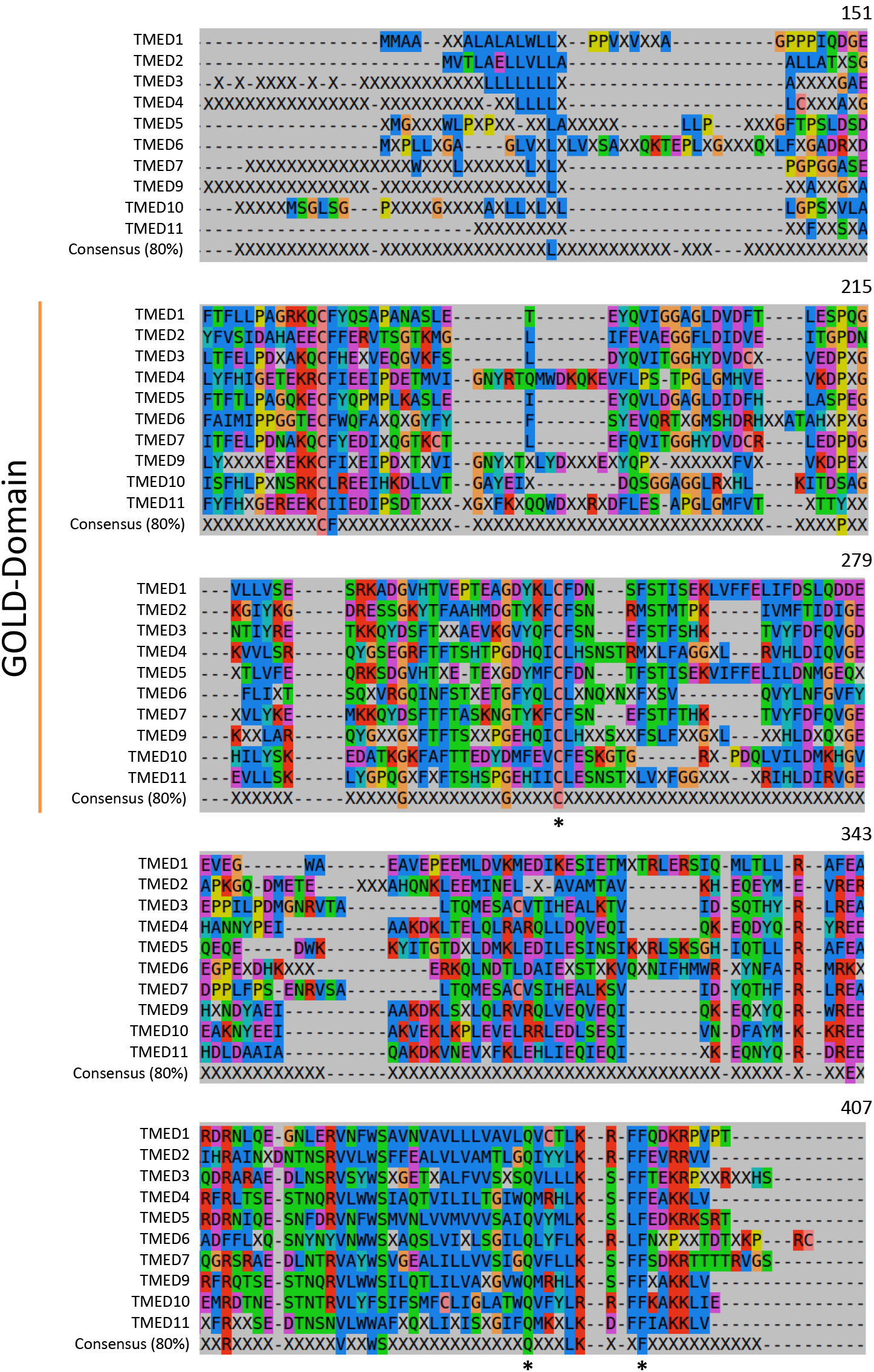
**A** Phylogenetic tree of TMED proteins. All mammalian sequences were collected from ENSEMBL and inaccurate sequence reads were deleted – 326 sequences were aligned by Clustal and presented in Newick format, actual distances are indicated in the graphical representation. TMED7 is close to TMED2 and comparison of 15 residues involved in hydrogen bonding to carbohydrate (or in close proximity) in TMED2 shows that 33 % (green) of these are conserved/conservatively replaced in TMED7, out of 23 residues implicated in Wnt binding 35 % (red) are conserved in TMED7 (identity), corresponding numbers are indicated for the other proteins; these show a higher substitution rate. **B** Clustal multiple-sequence alignment of mammalian (ENSEMBL) TMED sequences. 290 sequences were selected that did not contain gaps within the GOLD-domain (due to inaccurate sequence reads or splicing). The 80 % consensus of each TMED protein and the family consensus are presented. TMED4, 9, 10 and 11 contain inserts within the GOLD-domain (indicated). X represents the noconsensus position. Few amino acids are preserved in the whole family at the 80 % level, these accumulate at the transmembrane junction and within the transmembrane span, two, the Q and F (indicated with an asterisk) have been described as traffic signals (17, 20), the C within the GOLD-domain may be implicated in ligand binding.

**Figure 5.**
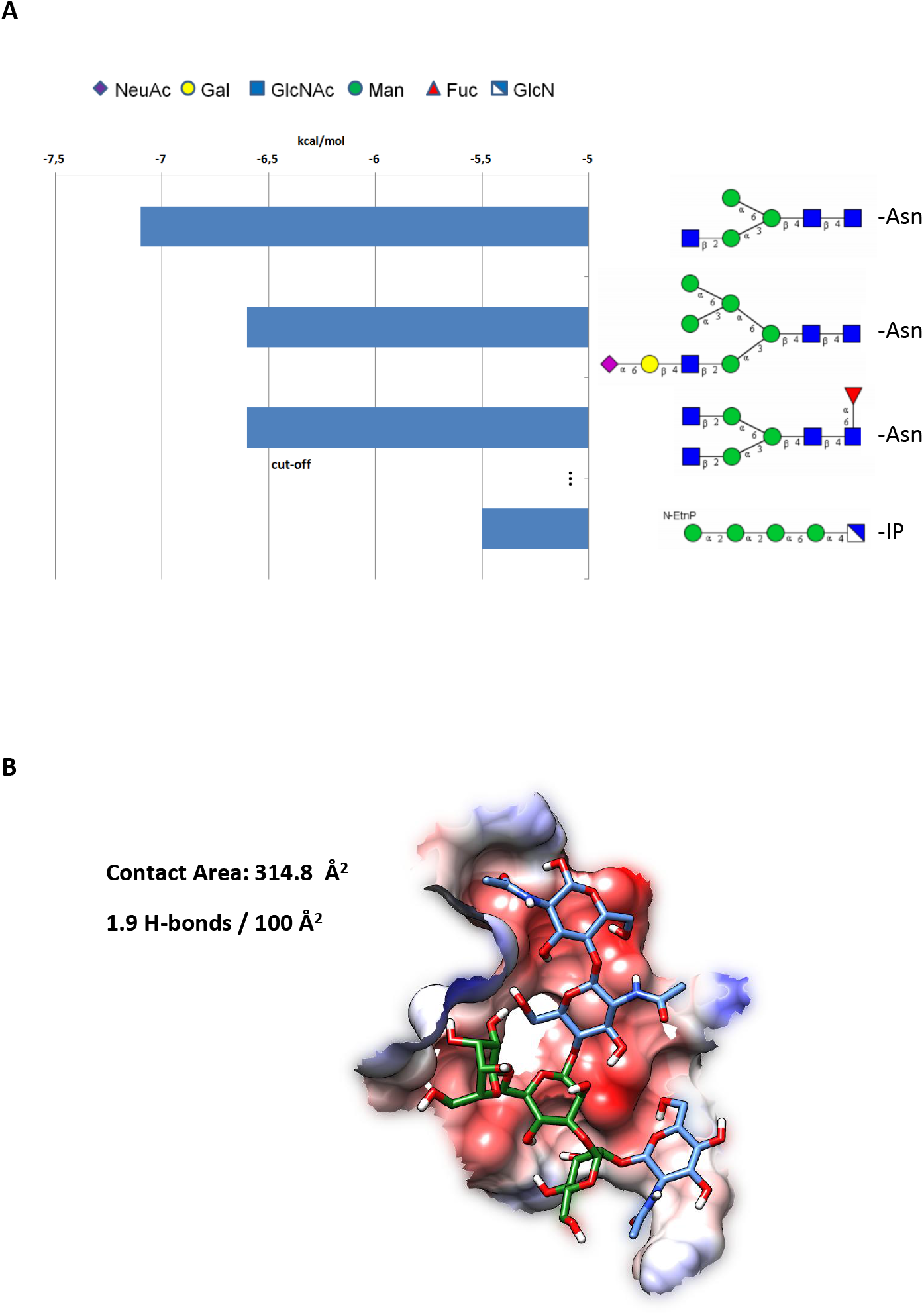
Docking of glycans. **A** N-glycans of complex, hybrid and high-mannose type as well as some terminal glycan groups were docked to p24 (GOLD-domain). The HybridM3 glycan represented a top score, further hybrid and complex glycans scored lower (see right (IUPAC code))(see Suppl. Tables S3/4). The core of glycosylphosphatidylinositol-anchors was included for comparison. Asn- and phosphati-dylinositol linkages (IP) are indicated. **B** Contact area of the N-glycan with p24-GOLD. The A-D-Manp- (1–6)-[B-D-GlcpNAc-(1–2)-A-D-Manp-(1–3)]-B-D-Manp-(1–4)-B-D-GlcpNAc-(1–4)-B-D-GlcpNAc-OH hybrid N-glycan with highest energy of interaction displays a contact area of 315 Å^2^ at 1 Å Van der Waals distance and 1.3-1.9 H-bonds per 100 Å^2^ contact area. The presentation is colored in Coulombic surface colors (see Fig. 2). Glycan monosaccharides are filled with IUPAC coloring.

The contact area of the GOLD-domain displays a surface of 314.8 Å^2^ with ~1.9 hydrogen bonds per 100 Å^2^ (Fig. 5B). Surprisingly, this seems very similar in configuration and extent to the hydrogen bonding capacity of β-sandwich type B lectins of the carbohydrate module (60) (CBM) clan. These are listed next to β-trefoil, cysteine knot or hevein folds forming 39 families of CBMs that are important in kingdoms from bacteria and plants to animals (60) (Suppl. Table S2). The carbohydrate affinity would be steered by hydrogen bond formation using backbone- and side-chain coordination from residues Arg69 at the N-glycan asparagine-linked terminus followed by additional bonding with Gly67, Asp68 and presumably Lys86 (Fig. 3A). The N-glycan (reducing) terminus forms two hydrogen bonds in the docking procedure. In molecular mechanics simulation the hydrogen-bonding network is extended to 5 hydrogen bonds and includes additional bonds to Asp53 and Glu55. It is likely that the two bonds to Arg69 would not be saturated since often the asparagine-linked GlcNAc is coordinated by hydrogen bonding to the covalently bound host protein.

### Docking of p24-GOLD

Since the binding site showed a concave face likely similar to a collagen-binding domain, I also scrutinized the putative collagen affinity by a hard-sphere docking method (49). Surprisingly, the affinity of p24-TMED2 to a collagen-like protein, that had been structurally determined and had previously been used in docking analyses (59), proved slightly superior to the interaction of a collagen-like polypeptide with the bacterial collagen-binding domain from *Clostridium histolyticum* itself (see Table S3). The models that were selected by the docking procedures with scores of ~7729 (see Table S1) with the disulfide rotameric form (Cys27 and Cys88) showed the preferential interaction at the N-termini of the collagen. Contrarily, top scores of models with energy minimized side-chains at residues 27/88, which according to the Dunbrack library (67) had probabilities of 0.057 (Cys27) and 0. 238 (Cys88) in reduced form, showed preferential interaction with the C-termini of the collagen-like polypeptide (Fig. 3C). To further explore the unique collagen interaction of the possibly promiscuous GOLD-domain, the PDB archive was matched to the UniProt database to obtain the 265 unique secreted proteins that had been determined at high structural resolution (Fig. 3D). Entries were selected and further scrutinized by aforementioned docking method to result in 15 highly scoring and optimized docked complexes. Among these I found the amyloid-precursor protein Kunitz-protease inhibitor domain scoring with −4.8 kcal/mol.

With respect to further complexes I established the high-affinity of the collagen-TMED2-GOLD model with −10.6 kcal/mol with the energy minimized protein (RMSD 0.245); on top of the collagen interaction, 1DZG, an antithrombin variant was highly scoring with −10.1 kcal/mol suggesting that additional protease inhibitors and domains may be functionally bound by TMED proteins. It is evident from the analyses of docked complexes that none of the other docked factors largely overlaps in binding relative to the docked collagen interface and that the collagen and glycan interface would exclusively count 93 % identical residues (Suppl. Fig. S5). Carbohydrate affinities could not be listed with docked factors since these could not be determined with PDBePISA (Fig. 3E), but are qualitatively similar to collagen interaction. Current limitations in docking for collagens include, furthermore, the non-flexible protein procedures.

GPI-anchors did not show considered binding affinity in this screen and high-affinity interaction of GPI-anchors of CD59 or Thy1 was so far not established with the GOLD-domain (Fig. 5A). The extended GPI-anchors of CD59 and Thy1 scored lower than the GPI-core with −5.5 kcal/mol (see Suppl. Table S4). A GPI-anchor interaction cannot be excluded, however, interactions with energies higher than −6.0 kcal/mol were not obtained with all studied conformers of TMED2-GOLD shown here.

TMED proteins have often been described as chaperones that may participate in protein folding in the secretory pathway. Machinery of the protein quality control was lately found also in the cis-/medial-Golgi and it is plausible that specific chaperone functions may be established (68). In this striking example of consistent biochemical (69) and structure evidence I show here, the docking of GOLD-domain residues 16-109 of p24 to the thumb domain of Wnt8. TMED2 is similarly docked to Wnt as Fz (Frizzled)(Fig. 3B). The palmitoleoyl group would be distant from the p24-Wnt binding interface, yet, the index finger interactions entirely cover, and possibly block Wnt-Fz binding with the ΔG=-18.3 kcal/mol (p24) versus ΔG=-6.4 kcal/mol (Fz cysteine-rich domain)(see Suppl. Table S1). Residues in contact with Wnt8 (Fig. 3B) are indicated in TMED2 with the surface side-chains (Fig. 2); a 12 % overlap of the Wnt binding site to the collagen-binding surface can be found (see Fig. 3E). Wnt-Fz interaction includes additional energy of interaction that e.g. at the cell surface entails the palmitoleoyl group at the thumb of Wnt8 attaching to Fz. Hydrogen bonding with complex/hybrid carbohydrates to p24 may replace the Fz lipid/proteinaceous interactions and include the interface at the index finger to replace and suppress Wnt-Fz signaling in early stages of plasma membrane travel in the ER and Golgi complex. Just as coatomer, GRASPs (Golgi reassembly and stacking proteins) may also be involved in their itinerary since these have been demonstrated to interact with cytoplasmic domains of TMED2 and 9 (70). Other cargo receptors may also interact with TMED proteins since e.g. Surf4, a cargo receptor showing 5 transmembrane domains, was found interacting with TMED2 and 10 in a complex in the presence of calcium (71).

Although the complete structure of Wnt has not been determined and a stretch of ~60 amino acids of Wnt11 cannot be modeled, it is tempting to speculate that the terminal N-glycosylation at Asn40 (72) might contribute to Wnt11 and p24 interactions: none of the modeled Wnts of *Xenopus laevis* (representative Wnt1/2/3a/4/5b/6/7/9b/10/11/16 including Wnt8) shows N-linked carbohydrates with proper geometry (predicted with NetNGlyc and GlyProt) that would impede docked p24 interaction. Models 1-3 and 5 can principally be excluded due to N-glycosylation since the binding site may be covered (Fig. 3B) when an N-glycan would be interposed in between the p24 and Wnt-palm domains. Thus, all models except model no. 4 could then be deselected according to this procedure since all Wnts 1/3-16 can be, and some have been (Wnt11: Ans40, Ans90, Asn300 / predicted Asn40, Asn90, Asn300, Asn304 *(Xenopus laevis);* Wnt3a: Asn87, Asn298 / predicted Asn63, Asn272 *(Xenopus laevis);* Wnt1 (dWnt-1): Asn103 *(Drosophila melanogaster))* shown to be N-glycosylated (72, 73) (Suppl. Fig. S4). Wnt would tumble when approaching the p24 from other directions and orientations and, possibly, Wnt-palm glycan binding would precede stably locked Wnt positioned similar to the geometry shown in Figures 3 and S4. Given current structural modeling, Wnt8 itself does not show a core N-glycan that would productively interact with p24 in this locked configuration (Fig. 3).

### TMED family

The generality of the TMED2 carbohydrate interaction was tested by alignment of all 29 mammalian current ENSEMBL TMED2 sequences. This showed that all residues involved in hydrogen bonding to the hybrid-Man3-GlcNAc N-glycan (Fig. 3A) were conserved at the 80 % cut-off (Suppl. Fig. S5). Some species contained gaps in sequence reads within the GOLD-domain, so that given a full genome, consensus above 90 % could be reached. Clustering all mammalian TMED proteins demonstrates a phylogenetic relationship (Fig. 4A) that groups TMED2, TMED6 and TMED10 separate from all other TMED proteins. TMED1 and 5, TMED3 and 7 as well as TMEDs 4, 9 and 11 are more highly related within their respective sub-groups as previously alluded to by phylogenetic tree analysis (74). With respect to ubiquitous Wnt interaction of TMED proteins, the visual inspection of the Prosite sequences showed that 14 out of 19 amino acids within the TMED2 GOLD-domain interacting with Wnt8 (Suppl. Fig. S1; conformer A) are abundant in the Prosite Logo when counting the first three frequently found amino acids (Suppl. Fig. S6; all species). Clustering analysis shows that despite the conservation of few residues in the mammalian TMED family (Consensus 80%, Fig. 4B) one residue within binding distance from the docked N-glycan is conserved in the entire family (cysteine)(labeled with *). The exact number of conserved residues in N-glycan binding (H-bonds) or Wnt8 interaction are indicated in Figure 4A (green, carbohydrate; red, Wnt8). This shows that among TMED proteins TMED7 and TMED3 are most highly related to the model of TMED2 with 33 % / 35 % and 27 % / 26 % conserved residues in hydrogen-bonding and Wnt interaction, respectively.

Further inspection shows, that also N-glycans of TMED proteins shown to be modified by carbohydrates within the GOLD-domain, had already been further analyzed (65, 75). For TMED7 and TMED9, N-glycans processed to complex and high mannose/complex N-glycans, respectively, had been found. Comparison of N-glycosylation sequons of each TMED of *Mus musculus* suggests that TMED4 and TMED11 could be N-glycosylated within, and likely TMED6 and 10 exterior to the GOLD-domain putative glycan binding pocket (data not shown). Asn103 in TMED7 is not located within the binding site of complex/hybrid N-glycans as gleaned from the structural comparative sequence analysis (Fig. 3A, Fig. 4B and Suppl. S5). The three TMED proteins TMED4, TMED9 and TMED11 may thus be impeded in putative glycan binding to the concave lectin surface if themselves covalently glycosylated within the GOLD-domain. It is possible that all other TMED proteins are free to interact with ligands via their concave patch GOLD-domain without steric hindrance.

## Discussion

### Wnt pathway

The Wnt pathway is considered as the key initiating mechanism involved in the development of colorectal cancer. Among other proteins, Wnts and collagen have been shown to be involved in epithelial-mesenchymal transition (EMT) (34,76,77). The APC (adenomatous polyposis coli) protein is part of a destruction complex, is involved in degradation of multiple factors including β-catenins and prevents their nuclear localization and TCF/LEF (T-cell transcription factor/lymphoid enhancer binding factor) signaling and is itself inactivated by somatic mutations. TCF/LEF triggers transcription of its target genes such as AXIN2, cyclin D1, Myc, SOX9, MMP7, BMP4, VEGF, EPHB, Dickkopf1 and TIAM1. Wnt secretion requires a multi-spanning cargo receptor, Wntless/Evi (evenness interrupted), shown to steer the Wingless-dependent patterning processes in *Drosophila melanogaster.* The cargo receptor Wntless/Evi was found to be essential in donor cells of the pathway but not on Wnt signaling receptor tissue (78, 79). Importantly, the Wnt-Tcf pathway was found to be modulated by members of the TMED/p24-family of cargo receptors: A screen for suppressors of metastasis has shown that among several genes only knockdown of TMED3 detrimentally affected growth of human cancer cells in mouse lung metastasis and led to cell spreading in hanging drop-aggregate tissue culture instead of to compaction of clonospheres (80). It was inferred that SOX12 and TMED3, involved in transcriptional regulation or contrarily in protein secretion, both affect the Wnt-TCF pathway. Interestingly, collagen binding to TMED2 could affect the secretion of extracellular matrix components that are involved in EMT. Boutros and colleagues had demonstrated that TMED5 *(opossum,* p24γ2) interacts with Wnt1 in *Drosophila melanogaster* and genetic interactions had also been shown for TMED2 (CHOp24, p24β1) – Wnt1 and in between TMED5 – WntD (Wnt8)(69). These results are consistent with the findings in human colorectal cancer with respect to Wnt localization and secretion and presumably altered growth and patterning. Also TMED4/9 *(eclair)* and TMED10 *(baiser)* had previously been found to affect Dpp (decapentaplegic) signaling and TMED function maintained the activity of maternally expressed type I TGF (transforming growth factor)-β receptor (81). Thus multiple signaling pathways are directly or indirectly affected by cargo receptor delivery.

### Early traffic in the secretory pathway and cargo receptors

The p24 protein may, as shown in this structural modeling, directly mediate transport or chaperone the Wnt-p24 complex to destined secretion and release at the plasma membrane. CD59, a GPI (glycosylphosphatidylinositol)-anchored protein, interacts with TMED2 and is guided to the plasma membrane by lipid rafts (82), thus lending credence to the idea that the early secretory pathway including the cis-/medial-Golgi is involved in apical delivery. Yeast cells, just as cells of mammalian origin, involve secretion with early control stations that trigger and regulate not only polarized but general exit (24, 83). Although the GPI-anchor interaction with TMED2 shown here is of intermediate affinity and interactions have been ascribed to the membrane proximal α-helical region in TMED5 (p24γ2)-TMED1 (p24γ1) chimeric constructs (84), complex/hybrid N-glycan binding may aid in sorting of GPI-anchored proteins if these are additionally glycosylated. Neither the GPI-anchor displaying only a single ethanolamine group nor the GPI-anchors with two or three ethanolamines showed high binding affinity in this screen.

Wnt proteins, and in particular Wnt11, may display bidentate interaction with carbohydrate-and protein-protein interfaces with the TMED2 GOLD-domain. Within the collagen and carbohydrate binding site a single cysteine (Cys27) is found which likely is bound to the second intra-sheet cysteine. This disulfide bridge may form in the ER-Golgi apparatus interface but since protein disulfide isomerases are also found at the plasma membrane, it is plausible that redox equilibria exist that allow for activity of -SH groups (85) at later stations in the secretory pathway. Thiols that are free to interact could affect cargo binding and the TMED protein plasma membrane travel. The collagen interaction of TMED proteins that is predicted based on the structural similarity and docking affinity of approximately −10.6 kcal/mol is likely to occur within the concave face of TMEDs. Collagen interacts possibly in two different orientations with the Cys-or disulfide-forming rotamers of TMED2-GOLD and should be further scrutinized.

### Outlook

The TMED functions may extend from chaperoning proteins along the secretory pathway, and decoy-receptor binding as shown for TMED2-Wnt here, to early inhibition of proteases that are active in intra-membrane proteolysis (86). The present analysis had demonstrated that TMED2 likely binds to carbohydrates with an affinity only marginally lower than VIP36 which was screened against a library of 159 glycans (data not shown)(Δ = 0.6 kcal/mol). TMED7 proteins, which according to this analysis may be, next to TMED3, most similar in glycan interactions to TMED2, have previously been shown to enhance cell surface abundance of Toll-like receptor 4 (TLR4) (87). It will be interesting to decipher direct lectin roles of TMED7 in innate immunity. Any function of TMED2 in collagen traffic should be established in the future which may not least include synchronized expression with chondrocyte differentiation as shown for TMED10 (88).

## Acknowledgements

Dr. Elise Wattendorf is thanked for critically reading the manuscript.

## Authors’ contributions

KF conceived and carried out the study, drafted the manuscript and prepared tables, figures and graphics. The author read and approved the final manuscript.

## Note added post-submission 2015 to Research Gate / 2016 to bioRxiv

The recent publication of Nagae et al. [Nagae M, Hirata T, Morita-Matsumoto K, Theiler R, Fujita M, Kinoshita T, et al. 3D Structure and Interaction of p24β and p24δ Golgi Dynamics Domains: Implication for p24 Complex Formation and Cargo Transport. J Mol Biol. 2016;428:4087–99] fully confirms the model of p24-TMED2 and shows that p24β1 and p24δ1GOLD domains share a common β-sandwich fold and may suggest, that in dimeric p24 proteins, the intra-sheet cysteines found could promote its dimerization or be closed in the (intramolecular) intra-sheet manner; if accessible to ligands, the dimeric binding site could provide a large β-sheet interaction that extends along the entire intermolecular sheet structure.

## Supplementary material

**Table S1.**
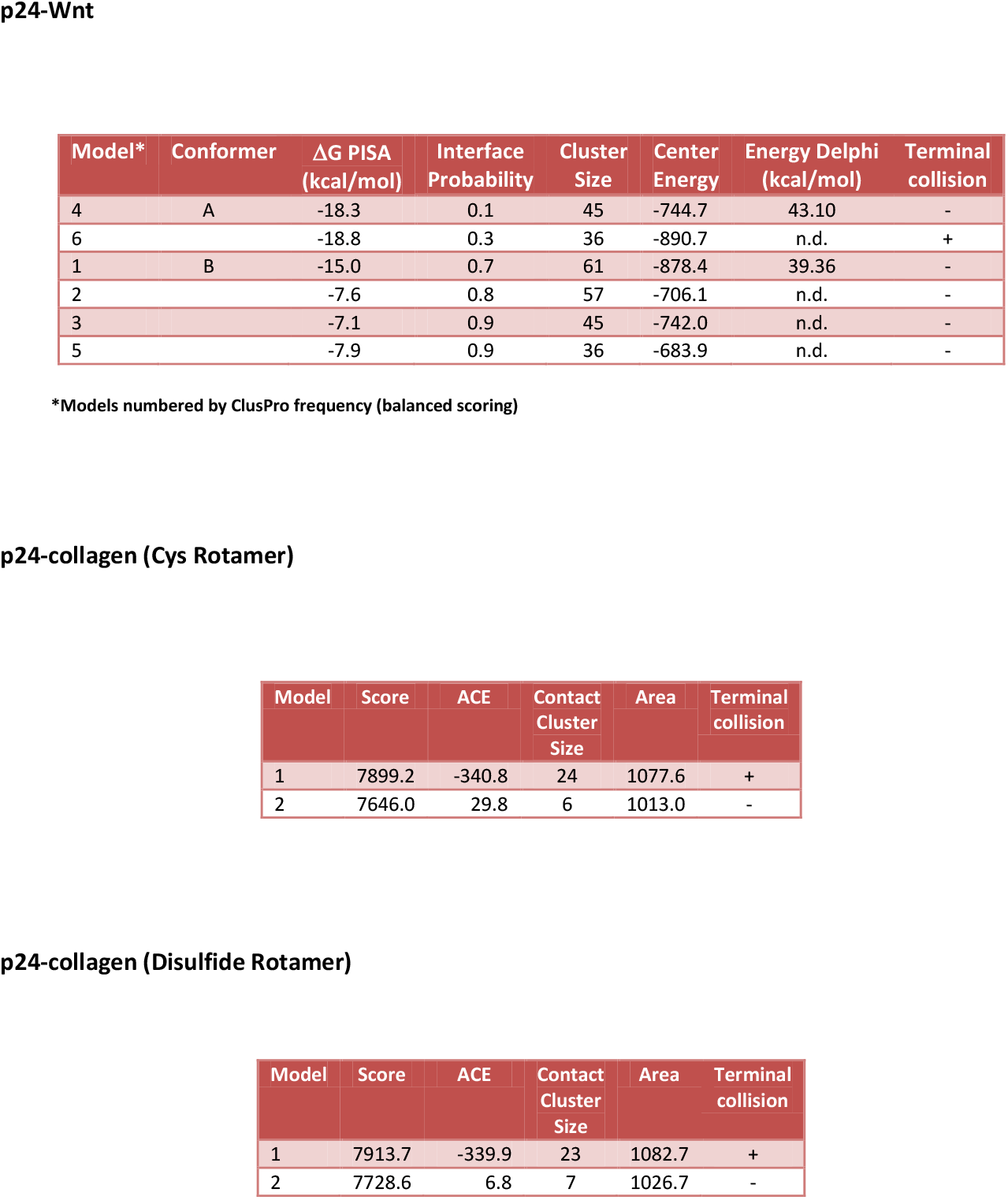
ClusPro, Patch- and FireDock, PDBePISA, DelPhi and structural analysis of GOLD-p24. **p24-Wnt**. TMED-Wnt models were ranked by interface probability. The ClusPro models of Wnt8-p24 complexes were analyzed for energy of interaction (ΔG PISA), PISA interface (interface probability), ClusPro Members (Cluster Size), ClusPro energy (center energy) and electrostatic energy of interaction (Energy DelPhi). Model numbering is according to ClusPro frequency, two conformers were further analyzed (A, B). **p24-collagen (Cys-Rotamer) and p24-collagen (Disulfide Rotamer)**. Geometric docking by PatchDock was used in further procedures; the PatchDock results with collagen which could be exclusively docked with the geometric algorithm are shown with the Cys-rotamer and disulfide-rotamer form indicated. Score is according to the average output (Score) for all docked poses in a contact cluster (defined by geometric point of interaction). Average atomic contact energy (ACE), cluster size (Contact Cluster Size) and average area of interaction (Area) are also indicated. All complexes were visualized and inspected, model no. 6 (Wnt8) and models 1 each (collagen cys-/disulfide-rotamer) showed a collision of p24 C-termini with the wnt and collagen proteins and can be excluded.

| Model* | Conformer | $\Delta G$ PISA<br>(kcal/mol) | Interface<br>Probability | Cluster<br>Size | Center<br>Energy | Energy Delphi<br>(kcal/mol) | Terminal<br>collision |
| --- | --- | --- | --- | --- | --- | --- | --- |
| 4 | A | -18.3 | 0.1 | 45 | -744.7 | 43.10 | - |
| 6 |  | -18.8 | 0.3 | 36 | -890.7 | n.d. | + |
| 1 | B | -15.0 | 0.7 | 61 | -878.4 | 39.36 | - |
| 2 |  | -7.6 | 0.8 | 57 | -706.1 | n.d. | - |
| 3 |  | -7.1 | 0.9 | 45 | -742.0 | n.d. | - |
| 5 |  | -7.9 | 0.9 | 36 | -683.9 | n.d. | - |

\*Models numbered by ClusPro frequency (balanced scoring)

**p24-collagen (Cys Rotamer)**
| Model | Score | ACE | Contact<br>Cluster<br>Size | Area | Terminal<br>collision |
| --- | --- | --- | --- | --- | --- |
| 1 | 7899.2 | -340.8 | 24 | 1077.6 | + |
| 2 | 7646.0 | 29.8 | 6 | 1013.0 | - |

**p24-collagen (Disulfide Rotamer)**
| Model | Score | ACE | Contact<br>Cluster<br>Size | Area | Terminal<br>collision |
| --- | --- | --- | --- | --- | --- |
| 1 | 7913.7 | -339.9 | 23 | 1082.7 | + |
| 2 | 7728.6 | 6.8 | 7 | 1026.7 | - |

**Table S2.** PDBeFold. The programme was used to search for structural homologues of the Golgi dynamics domain of Sec14-like 3 (residues 280-385). The output lists protein names (Protein), the Q-Score of structural similarity (Q-Score), the RMSD (RMSD), ligand (Ligand) and species (Species). The first three top scores display interaction with collagen on a concave interface (Collagenase collagen-binding domain) or with carbohydrate (Rets (Regulator of exopolysaccharide and type III Secretion) sensor domain and Endo-1,4-β-galactosidase). Multiple carbohydrate binding modules (CBM) with a typical jelly-roll fold are further scored. Interactions of carbohydrates with the jelly-roll occurs at multiple locations, GOLD-domains of p24 would according to this analysis be classified similar to B-type CBMs. Galectins may be included but show extended sheet structures with an unequal number of single β-strands.

| Structure Alignment Results: Human Sec14-like 3 (modular Golgi dynamics domain 280-385) |  |  |  |  |
| --- | --- | --- | --- | --- |
| Protein<sup>#</sup> | Q-Score | RMSD | Ligand | Species |
| Collagenase collagen-binding domain | 0.42 | 2.06 | collagen | Clostridium histolyticum |
| Rets sensor domain | 0.31 | 2.42 | carbohydrate | Pseudomonas aeruginosa |
| Endo-1,4-β-galactosidase | 0.30 | 2.59 | carbohydrate | Thermotoga maritima |
| ... | ... | ... | ... | ... |
| XG-34 CBM | 0.29 | 2.76 | carbohydrate | Rhodotermus marinus |
| CBM36 | 0.29 | 3.21 | carbohydrate | Paenibacillus polymyxa |
| ... | ... | ... | ... | ... |
| Galectin 7 | 0.26 | 2.49 | carbohydrate | Homo sapiens |
| ... | ... | ... | ... | ... |
| Galectin 9 | 0.25 | 2.78 | carbohydrate | Homo sapiens |

\*Results were collated and identical structures were deleted

**Table S3.** **A** Docking of glycans. N-glycans of complex, hybrid and high-mannose type as well as some terminal glycan groups were docked to p24 (GOLD-domain) with docking parameters: grid size 40×46×35 in global and grid size 25×25×25 in local docking. The HybridM3 glycan represented the top score, additional hybrid and complex glycans scored lower. In exhaustiveness setting 11 (versus 8) no higher scoring was found for these entries. The core of glycosylphosphatidylinositol-anchors (GPI-core) was included for comparison. Result number 3, the complexM3-Fuc-GlcNAc-GlcNAc glycan, shows an altered core-GlcNAc configuration, all listed results bound to the described binding groove with high affinity in the global docking approach. **B** Collagen was analyzed for docking to p24-GOLD by Patch-Dock. Energies of docking were determined by PDBePISA to allow comparison with ClusPro PDBePISA values - these scored significantly lower than FireDock values but generated plausible values with both, the TMED2 cys-rotamer and disulfide-rotamer. RMSDs of the optimized proteins were (2) 0.247/0.224/0.150/0.165 and (4) 0.245/0.211/0.143/0.162 for TMED2 and collagen chains A-C. Control docking results with a bacterial collagen-binding domain from *Clostridium* showed values from – 6.7 to -9.9 kcal/mol of PDBePISA interaction in a docking configuration to 1NQD (calcium-bound) that was similar to a previously analyzed model (59). This was close to TMED docking energies. Energies of interaction with calcium were neglected for 1NQD and drive the interaction towards binding of both, calcium and collagen but do not contribute to collagen interaction as such. Top entries of the Patch-Dock cluster were energy-optimized by Chimera energy-minimization (combined models) and yielded elevated energies of interactions (opt).

| No. | Structure* | Energy<br>(kcal/mol) |
| --- | --- | --- |
| 1 | HybridM3-GlcNAc | -7.1 |
| 2 | HybridM5-GlcNAc-Gal-NeuNAc | -6.6 |
| 3 | ComplexM3-Fuc-GlcNAc-GlcNAc** | -6.6 |
| ... | ... | ... |
| 24 | GPI-core | -5.5 |

| Run | Structure* | $\Delta G$ PISA<br>(kcal/mol) |
| --- | --- | --- |
| 1 | Cys-Rotamer | -9.9 |
| 2 | Cys-Rotamer opt. | -11.1 |
| 3 | Disulfide | -9.3 |
| 4 | Disulfide opt.<sup>#</sup> | -10.6 |

\*PatchDock top scoring molecules of model 2 each (from collated contact cluster) were used non-optimized or structurally optimized (opt). <sup>#</sup>Cys27-Cys88 was linked by disulfide before optimization. The Cys- and disulfide-rotamer showed inverse orientations of the collagen with respect to N- and C-termini.

**Table S4.**
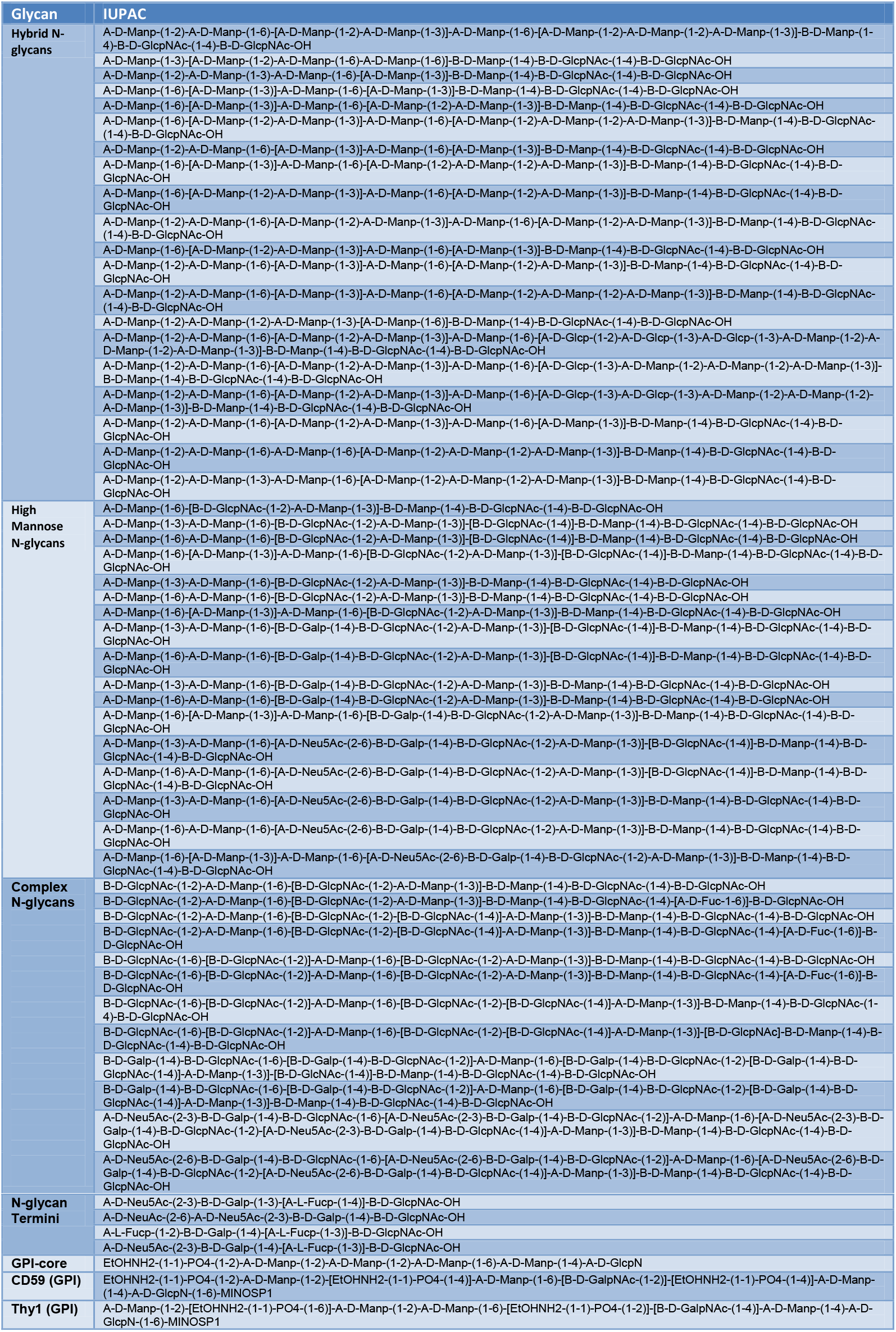
Glycans used in the screen. List of Hybrid N-glycans, High Mannose N-glycans, Complex N-glycans and N-glycan Termini (IUPAC). Three GPI-cores are included: the basic GPI-core, the CD59 GPI-anchor and the Thy1 GPI-anchor both with added MINOS (myoinositol)-phosphate.

|  |  |
| --- | --- |
| N-glycan<br>Termini | A-D-Neu5Ac-(2-3)-B-D-Galp-(1-3)-[A-L-Fucp-(1-4)]-B-D-GlcpNAc-OH |
|  | A-D-NeuAc-(2-6)-A-D-Neu5Ac-(2-3)-B-D-Galp-(1-4)-B-D-GlcpNAc-OH |
|  | A-L-Fucp-(1-2)-B-D-Galp-(1-4)-[A-L-Fucp-(1-3)]-B-D-GlcpNAc-OH |
|  | A-D-Neu5Ac-(2-3)-B-D-Galp-(1-4)-[A-L-Fucp-(1-3)]-B-D-GlcpNAc-OH |
| GPI-core | EtOHNH2-(1-1)-PO4-(1-2)-A-D-Manp-(1-2)-A-D-Manp-(1-2)-A-D-Manp-(1-6)-A-D-Manp-(1-4)-A-D-GlcpN |
| CD59 (GPI) | EtOHNH2-(1-1)-PO4-(1-2)-A-D-Manp-(1-2)-[EtOHNH2-(1-1)-PO4-(1-4)]-A-D-Manp-(1-6)-[B-D-GalpNAc-(1-2)]-[EtOHNH2-(1-1)-PO4-(1-4)]-A-D-Manp-(1-4)-A-D-GlcpN-(1-6)-MINOSP1 |
| Thy1 (GPI) | A-D-Manp-(1-2)-[EtOHNH2-(1-1)-PO4-(1-6)]-A-D-Manp-(1-2)-A-D-Manp-(1-6)-[EtOHNH2-(1-1)-PO4-(1-2)]-[B-D-GalpNAc-(1-4)]-A-D-Manp-(1-4)-A-D-GlcpN-(1-6)-MINOSP1 |

**Figure S1.**
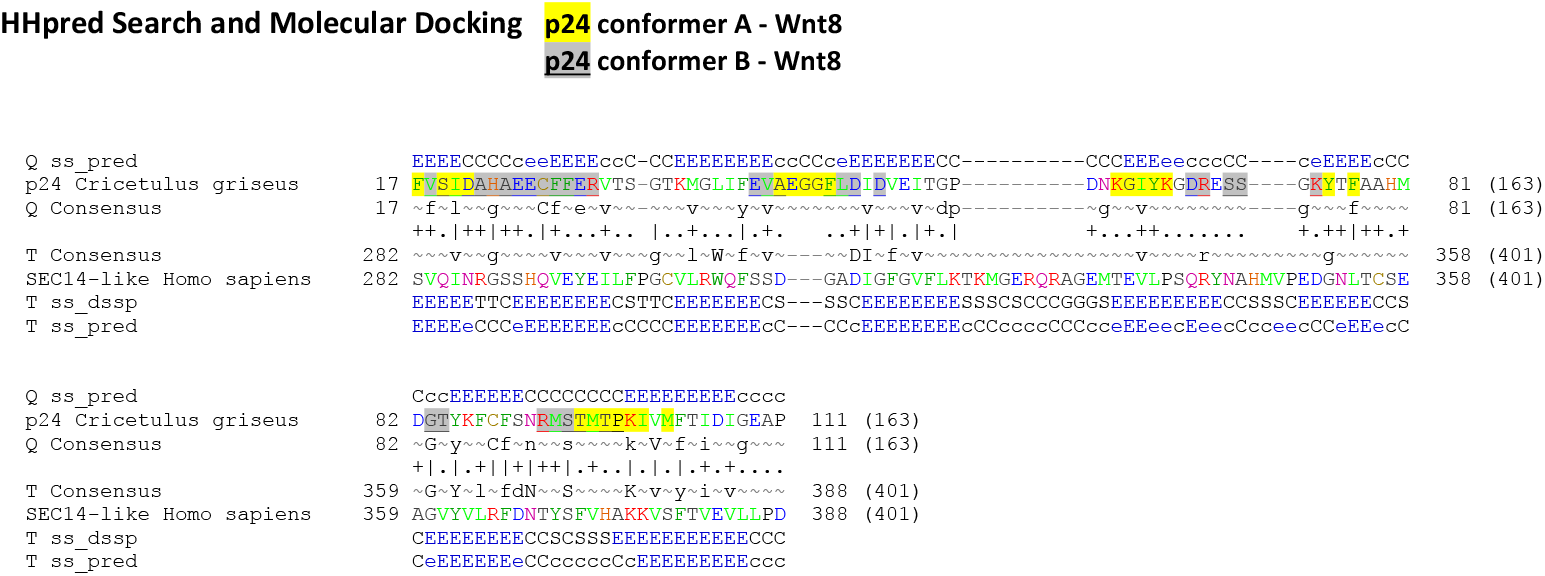
The HHpred search was used to search for p24 *(Cricetulus griseus)* homologues in the database (pdb). Upon iterative Psi Blasting, the secondary structure enhanced output (PsiPred) listed 100 factors none of which was superior to 0.016 in E-value except number 1, the Sec14-like protein 3, which scored with an E-value of 4.9^-10^, P-value of 1.4^-14^, and a score of 89.9. The overall probability of a true positive was 99.1%. 4UYB, the entry of the structure of the Sec14-like protein 3 *(Homo sapiens),* was aligned from positions 282-388 with the full length GOLD-domain of p24 (residues 17-111). The alignment generated by the procedure lists the secondary structure propensities (E/C) (ss_pred). Template DSSP values were calculated from the 3-D structures determined by X-ray crystallography or NMR (ss_dssp) and are indicated. The consensus sequences of the query (p24) or template (4UYB_A) is shown from multiple sequence alignment (Consensus). Molecular docking results from ClusPro with Wnt8 are indicated. Only residues with >20 % buried surface area are labeled within the amino acid sequence for conformers A in yellow and B in grey underlined, or in yellow underlined if present in both (A and B).

**Figure S2.**
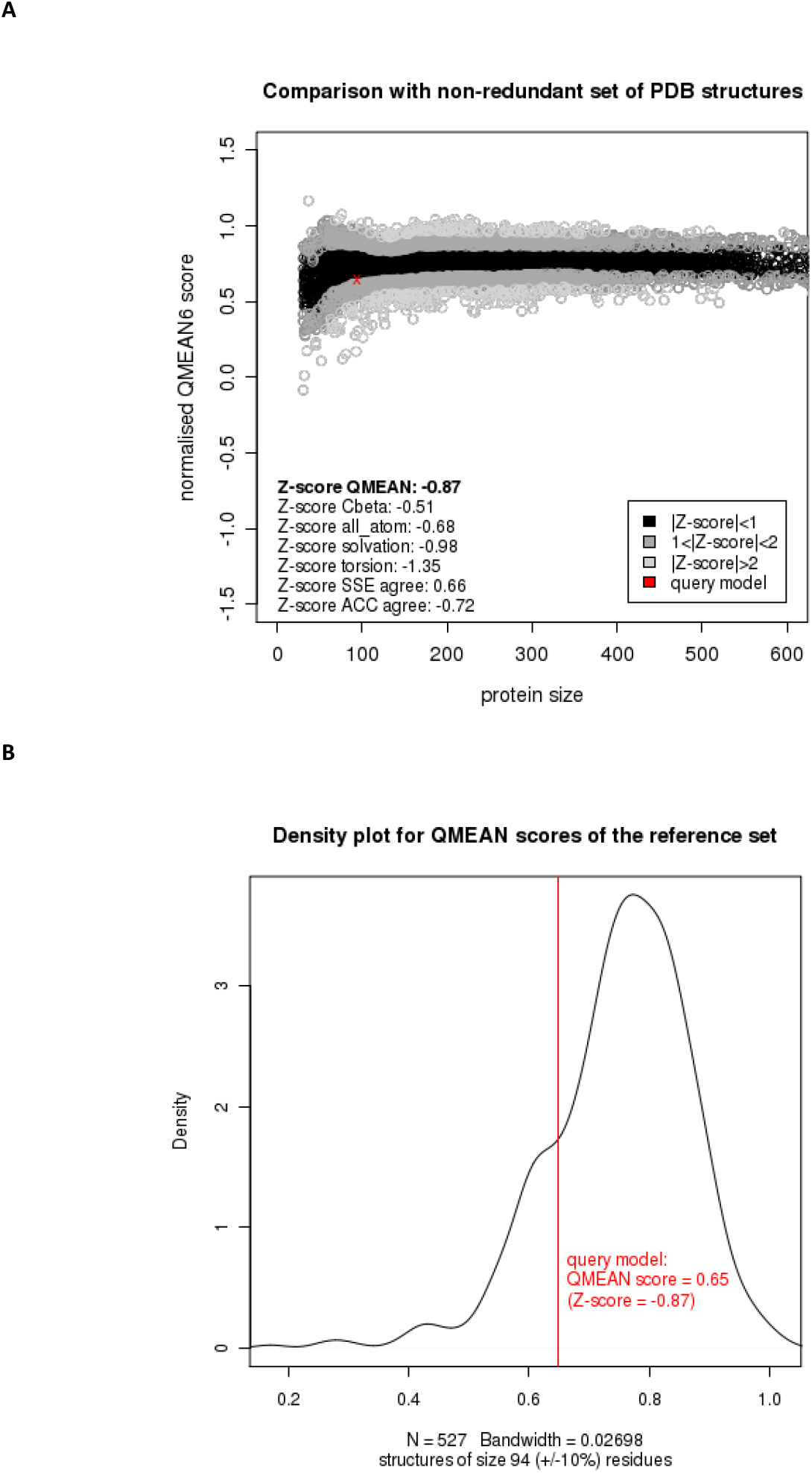
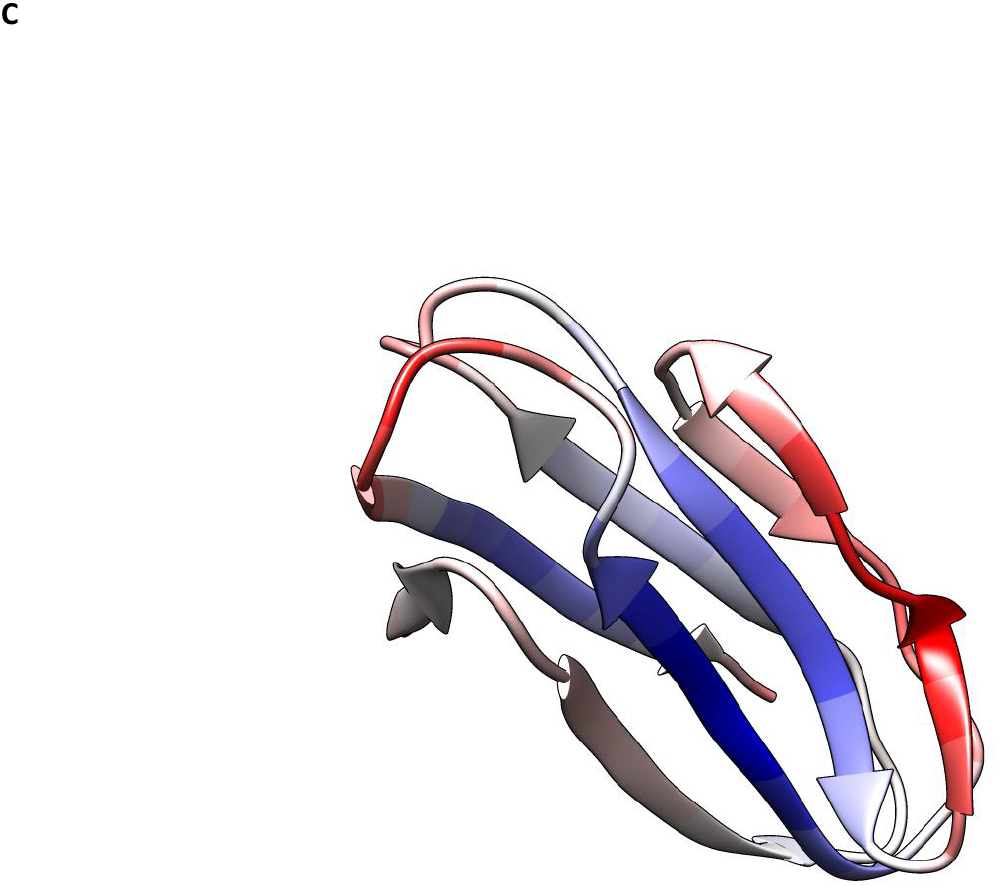
A QMean results. Comparison to the non-redundant set of PDB structures. Z-scores < ±1 are indicated in black, Z-scores < ±2 in grey, Z-scores>±2 in light grey and the query model of p24 as a cross. These results were obtained from Cβ interaction (Cbeta), all-atom interaction (all_atom), solvation, torsion, predicted and calculated secondary structure- (SSE) and predicted-relative and relative solvent ac-cessibility-agreement (ACC) comparisons with indicated results (QMEAN6). B The Z-score and QMean score relative to structures of similar size; the distribution lists structures of 94 amino acid sequence length (±10%). The QMean score of 0.65 indicates a “good” model indicated with a red line (40). C Local B-factors. Thresholds were set to 0 (dark blue) and 5.1 (red). The GOLD-domain is shown from residue 16-109. The center of the binding groove shows a particularly low B-factor value. Orientation of the molecule is shown similarly to in Fig. 3.

**Figure S3.**
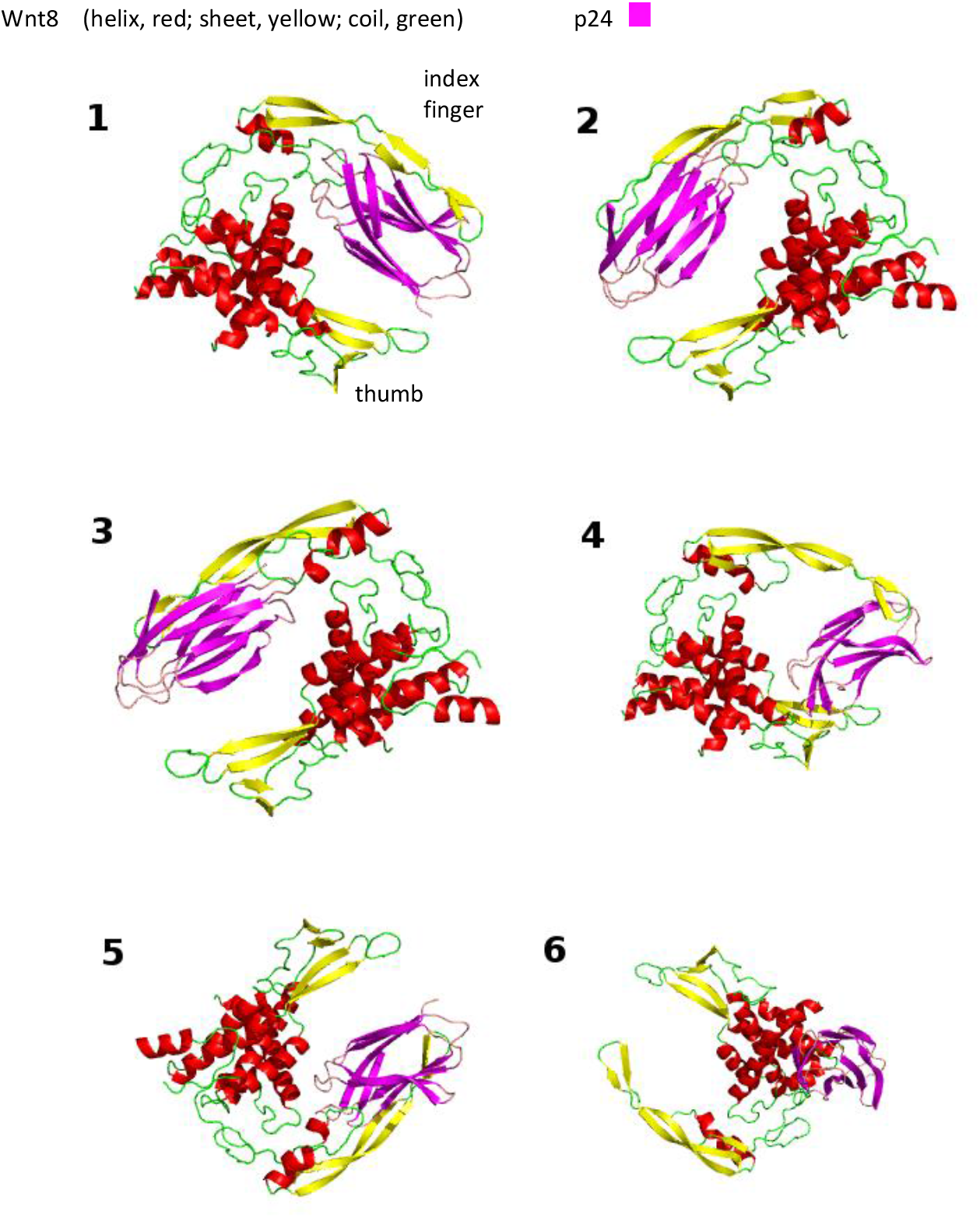
ClusPro results. Models obtained from the ClusPro queue are listed with score (1–6) from ClusPro balanced scoring (electrostatic, hydrophobic, van der Waals) and ranked according to cluster size (Table S1). The index finger, thumb of Wnt8 and p24 are indicated (see Fig. 3B).

**Figure S4.**
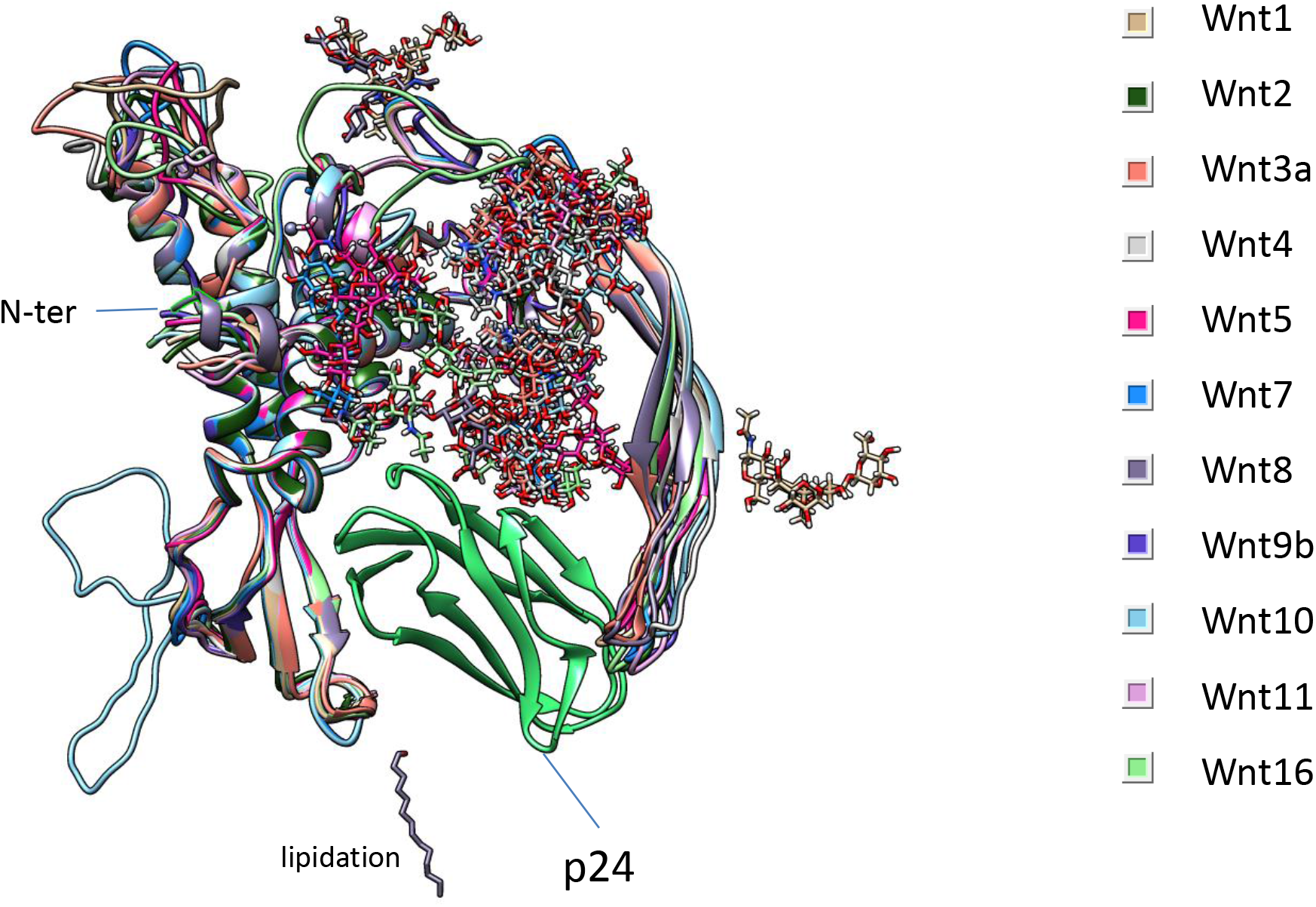
Modeled Wnt proteins. Wnt1, 2, 3a, 4, 5, 6, 7, 6, 9b, 10, 11 and 16 from *Xenopus laevis* were modeled with the 4F0A template (Wnt8)(color code). NetNGlyc was applied to predict N-glycosylation sequons and sites with scores above 0.5 were glycosylated with GlyProt (basic N-glycans, proper stereochemistry)(57). Wnt2 is not N-glycosylated. None of the added cores (GlcNAc-GlcNAc-Man3) impeded p24 binding in visual inspection and it is likely that even core extensions would not abolish p24-Wnt interactions. Plausibly, glycosylation may act as fishing bait to collect low-affinity cargo receptors to Wnts. Wnt6 was not modeled completely, would shift the binding site towards the left of the shown panel or not bind p24 in the given configuration, and has been omitted for legibility. The location of Wnt8 lipidation is indicated. The current result may suggest that acyl-chains do not interact with TMED in the shown fixed position due to a distance of ~20 Å from Ser187 (Wnt) to Ser90 (TMED) in the middle of the groove and a length of ~16 Å of an elongated acyl chain.

**Figure S5.**
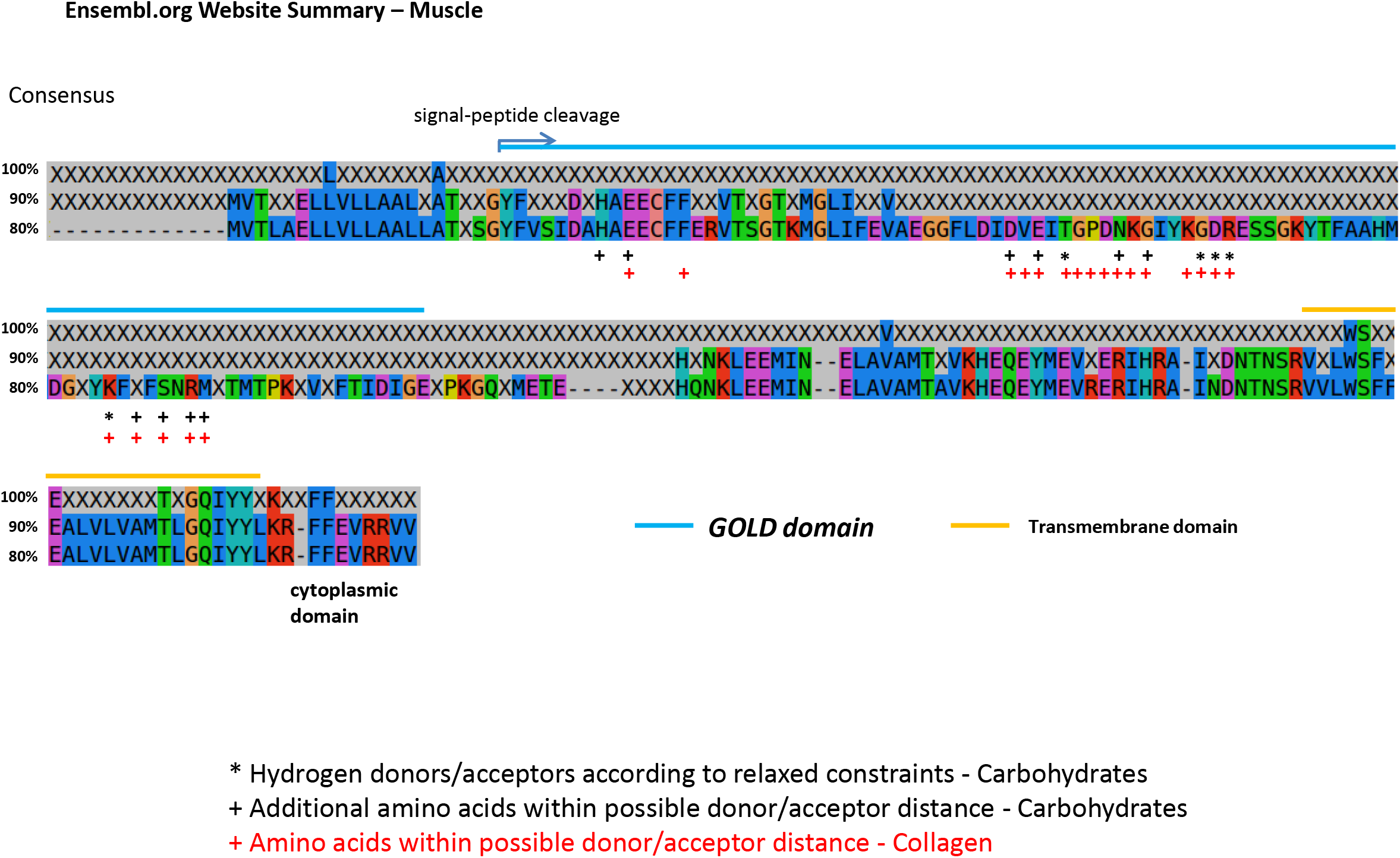
Display of TMED2 (p24ß1) members (full length), 29 mammalian sequences. Ensembl sequences were aligned, consensus sequences (80% – 100%) are shown. Amino acids are colored by properties. Hydrogen bond donors/acceptors (*) of glycans and additional amino acids within possible donor/acceptor distance (+) are indicated in black. Amino acids within proximity of docked collagen are indicated with red (+). These largely overlap with the glycan binding site. 93 % of residues of the glycan interface occur in both, the glycan interface and in the collagen binding site. The ENSFCAP sequence from cat was experimentally included, contains a short GOLD-domain from residues 16-63 (comparison to CHO TMED2), but is similar to a human splice variant of TMED2 (FLJ52154). Sequences included orthologues from Alpaca *(Vicugna pacos),* Armadillo *(Dasypus novemcinctus),* Bushbaby *(Otolemur garnettii),* Cat *(Felis catus),* Cow *(Bos taurus),* Dog *(Canis lupus familiaris),* Dolphin *(Tursiops truncatus),* Elephant *(Loxodonta africana),* Ferret *(Mustela putorius furo),* Gibbon *(Nomascus leucogenys),* Gorilla *(Gorilla gorilla gorilla),* Guinea Pig *(Cavia porcellus),* Hedgehog *(Erinaceus europaeus),* Human *(Homo sapiens),* Hyrax *(Procavia capensis),* Kangaroo rat *(Dipodomys ordii),* Lesser hedgehog tenrec *(Echinops telfairi),* Macaque *(Macaca mulatta),* Marmoset *(Callithrix jacchus),* Megabat *(Pteropus vampyrus),* Microbat *(Myotis lucifugus),* Mouse *(Mus musculus),* Olive baboon *(Papio anubis),* Orangutan *(Pongo abelii),* Panda *(Ailuropoda melanoleuca),* Pig (Sus *scrofa),* Pika *(Ochotona princeps),* Rabbit *(Oryctolagus cuniculus),* Rat *(Rattus norvegicus),* Sheep *(Ovis aries),* Shrew *(5orex araneus),* Sloth *(Choloepus hoffmanni),* Vervet *(Chlorocebus sabaeus)* and Wallaby *(Macropus eugenii).* The cleaved signal-sequence, the GOLD-domain (residues 16-109), the transmembrane and cytoplasmic domains are indicated.

**Figure S6.**
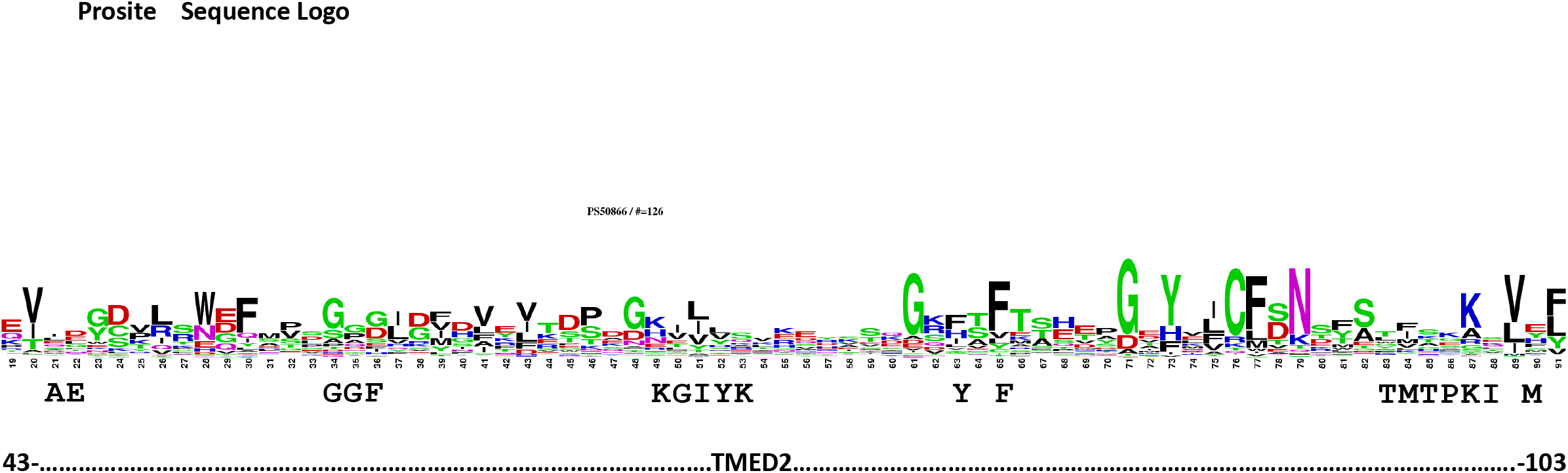
Prosite sequence Logo of the p24/Emp24/gp25L domain. 126 Prosite entries are represented in the sequence Logo. 14 of 19 amino acids interacting with Wnt8 (conformer A; see Fig. S1) will be identified as abundant if considered in the first three ranks of the Logo (more amino acids were determined to bind but are not entirely covered by the Logo). Amino acids of TMED2 are indicated in bold below the sequence Logo if interacting with Wnt8 and were annotated from visual CLUSTAL comparison (see Figure 2).

## Abbreviations

APC: adenomatous polyposis coli
CBM: carbohydrate binding module
CHO: chinese hamster ovary (from Cricetulus griseus)
Dpp: decapentaplegic
EMT: epithelial-mesenchymal transition
ER: endoplasmic reticulum
Evi: evenness interrupted
FRT: Fischer rat thyroid
Fz8: frizzled-8
GalNAc: N-acetylgalactosamine
GlcNAc: N-acetylglucosamine
GlcNAcT-I: GlcNAc-Transferase I
GOLD: Golgi-dynamics
Golgi-Man: Golgi-mannosidase
GPI-anchor: glycosylphosphatidylinositol-anchor
GRASP: Golgi reassembly and stacking protein
GSL: glycosphingolipid
LEF: lymphoid enhancer binding factor
Man: mannose
MDCK: Madin-Darby Canine Kidney
MINOS: myo-inositol
NeuAc: N-acetylneuraminic acid
PIG-A: PI (phosphatidylinositol) GlcNAcT gene
Rets: regulator of exopolysaccharide and type III secretion
TCF: T-cell transcription factor
TGF: transforming growth factor
TMED: transmembrane emp24 domain containing
TLR4: toll-like receptor 4
VIP36: vesicular-integral membrane protein of 36 kD
Wnt: wingless

